# Transient structural variations have strong effects on quantitative traits and reproductive isolation in fission yeast

**DOI:** 10.1101/047266

**Authors:** Daniel C. Jeffares, Clemency Jolly, Mimoza Hoti, Doug Speed, Liam Shaw, Charalampos Rallis, Francois Balloux, Christophe Dessimoz, Jürg Bähler, Fritz J. Sedlazeck

## Abstract

Large structural variations (SVs) in the genome are harder to identify than smaller genetic variants but may substantially contribute to phenotypic diversity and evolution. Here we analyze the effects of SVs on gene expression, quantitative traits, and intrinsic reproductive isolation in the yeast *Schizosaccharomyces pombe*. We establish a high-quality curated catalog of SVs in the genomes of a worldwide library of *S. pombe* strains, including duplications, deletions, inversions and translocations. We show that copy number variants (CNVs) frequently segregate within closely related clonal populations, are weakly linked to single nucleotide polymorphisms (SNPs), and show other genetic signals consistent with rapid turnover. These transient CNVs produce stoichiometric effects on gene expression both within and outside the duplicated regions. CNVs make substantial contributions to quantitative traits such as cell shape, cell growth under diverse conditions, sugar utilization in winemaking, whereas rearrangements are strongly associated with reproductive isolation. Collectively, these findings have broad implications for evolution and for our understanding of quantitative traits including complex human diseases.

---

A variety of genetic changes can influence the biology of species, including single-nucleotide polymorphisms (SNPs), small insertion-deletion events (indels), transposon insertions and large structural variations. Structural variations (SVs), including deletions, duplications, insertions, inversions and translocations, are the most difficult to type and consequently the least well described.

Nevertheless, it is clear that SVs have strong effects on various biological processes. Copy number variants (CNVs) in particular influence quantitative traits in microbes, plants and animals, including agriculturally important traits and a variety of human diseases^1–5^. Inversions are known to influence reproductive isolation^6–13^ and other evolutionary processes such as recombination^8^ and hybridization between species^14^, with a variety of consequences^15^.

We and others have recently begun to develop the fission yeast *Schizosaccharomyces pombe* as a model for population genomics and quantitative trait analysis^6,7,16–18^. This model organism combines the advantages of a small, well-annotated haploid genome^19^, abundant tools for genetic manipulation and high-throughput phenotyping^20^, and considerable resources of genome-scale and gene-centric data^21–23^.

Previous analyses of fission yeast have begun to describe both naturally occurring and engineered inversions and reciprocal translocations^6,7,18^. Given this evidence for SVs and their effects in this model species, we recognized that a systematic survey of SVs would progress our understanding of their biological influence. Here, we utilize the recent availability of 161 fission yeast genomes and extensive data on quantitative traits and reproductive isolation^17^ to describe the nature and effects of SVs in *S. pombe*.

We show that SVs have strong effects on various aspects of biology. They contribute an average of 11% of trait variance (the much more abundant SNPs contribute 24% on average), with the largest effects coming from CNVs. We show that CNVs are transient within clonal populations, and are frequently not well tagged by SNPs. We also show that rearrangements (inversions and translocations) contribute to reproductive isolation, whereas CNVs do not.

## RESULTS

### Genome- and population-wide detection of structural variations

To predict an initial set of SVs, we applied four inference software packages (Delly, Lumpy, Pindel and cn.MOPs)^24–27^ to existing short-read data^17^, using parameters optimized on simulated data (Methods). We then filtered these initial predictions, accepting SVs detected by at least two callers, to obtain 315 variant calls (141 deletions, 112 duplications, 26 inversions, 36 translocations). We release this pipeline as an open-source tool called SURVIVOR (Methods). To ensure a high specificity, we further filtered the 315 variants by removing SV calls whose breakpoints overlapped with low complexity regions or any that corresponded to previously annotated long terminal repeats (LTRs)^17^. Finally, we manually vetted all the remaining SVs by visual inspection of read alignments in multiple strains for all remaining candidates. This meticulous approach aimed to ensure a high quality call set, to mitigate against the high uncertainty associated with SV calling^25^.

This curation produced a set of 113 SVs, comprising 23 deletions, 64 duplications, 11 inversions and 15 translocations (Figure 1a). Reassuringly, when applying our variant calling methods to an engineered knockout strain, we correctly identified the known deletions and called no false positives. Attempts to validate all rearrangements by PCR and BLAST searches of *de novo* assemblies positively verified 76% of the rearrangements, leaving only a few PCR-intractable variants unverified (see Methods for details).

**Figure 1.**
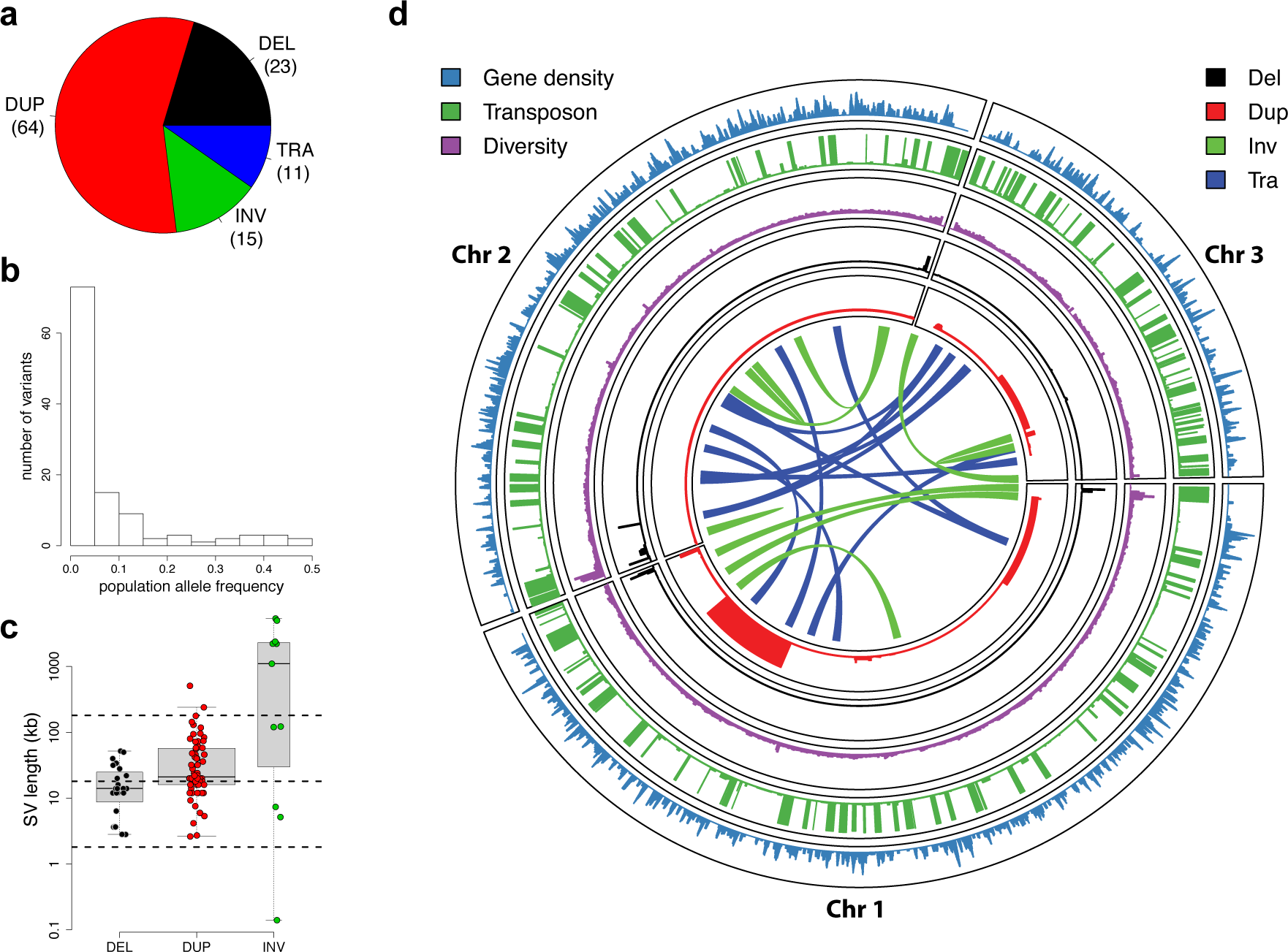
Characteristics of SVs in *S. pombe*. (**a**) Relative proportions of SVs identified. Duplications (DUP) were the most abundant SVs, followed by deletions (DEL), inversions (INV), and translocations (TRA). (**b**) Population allele frequency distribution of SVs, showing the frequencies of less abundant alleles in the population (minor allele frequencies). (**c**) Length distributions of SVs, log_10_ scale. Deletions were smallest (2.8–52 kb), duplications larger (2.6–510 kb), and inversions often even larger, spanning large portions of chromosomes (0.1 kb–5,374 kb, see (d)). Horizontal dotted lines show the size of chromosome regions that contain an average of 1, 10 and 100 genes in this yeast. (**d**) Locations of SVs on the three chromosomes compared to other genomic features. From outside: density of essential genes, locations of *Tf*-type retrotransposons, diversity (π, average pairwise diversity from SNPs), deletions (black), duplications (red), and breakpoints of inversions and translocations as curved lines inside the concentric circles (green and blue, respectively). Bar heights for retrotransposons, deletions and duplications are proportional to minor allele frequencies. Diversity and retrotransposon frequencies were calculated from 57 non-clonal strains as described by Jeffares, et al. ^17^.

Most SVs were present at low frequencies, with 28% discovered in only one of the strains analyzed (Figure 1b). The deletions were generally slightly smaller (median length 14 kb, Figure 1c) than duplications (median length of 21 kb), with the largest duplication extending to 510 kb and covering 200 genes (a singleton in strain JB1207/NBRC10570). The majority of CNVs were present in copy numbers varying between zero and sixteen (subsequently we refer to amplifications of two or more copies as ‘duplications’).

All SVs, particularly deletions and duplications, were biased towards the ends of chromosomes (Figure 1d, Supplementary Figures 1 and 2), which are characterized by high genetic diversity, frequent transposon insertions, and a paucity of essential gene^17^, similar to *Saccharomyces cerevisiae* and *S. paradoxus*^28,29^. All SVs preferentially occurred in positions of low gene density and were strongly under-enriched in essential genes (Supplementary Figure 2).

To describe SVs further, we conducted gene enrichment analysis with the AnGeLi tool (**Supplementary Table 1**), which interrogates gene lists for functional enrichments using multiple qualitative and quantitative lnformation sources^30^. The CN V-overlapping genes were enriched for caffeine/rapamycin induced genes and genes induced during meiosis (P = 4×10^−7^ and 1×10^−5^, respectively); they also showed lower relative RNA polymerase occupancy and were less likely to contain genes known to produce abnormal cell phenotypes (P = 1.8×10^−5^ and 3×10^−5^, respectively). These analyses are all broadly consistent with a paucity of CNVs in genes that encode essential mitotic functions. Rearrangements disrupted only a few genes and showed no significant enrichments.

### Duplications are transient within clonal populations

Our previous work identified 25 clusters of near-clonal strains, which differed by <150 SNPs within each cluster^17^. We expect that these clusters reflect either repeat depositions of strains differing only at few sites (e.g. mating-type variants of reference strains *h*^90^ and *h*^−^ differ by 14 SNPs) or natural populations of strains collected from the same location. Such ‘clonal populations’ reflect products of mitotic propagation from a very recent common ancestor, without any outbreeding. We therefore expected that SVs should be largely shared within these clonal populations.

Surprisingly, our genotype predictions indicated that most SVs present in clonal populations were segregating, i.e. were not fixed within the clonal population (68/95 SVs, 72%). Furthermore, we observed instances of the same SVs that were present in two or more different clonal populations that were not fixed within any clonal population. These SVs could be either incorrect allele calls in some strains, or alternatively, recent events that have emerged during mitotic propagation. To distinguish between these two scenarios, we re-examined the read coverage of all 49 CNVs present within at least one clonal population. Since translocations and inversions were more challenging to accurately genotype, we did not re-examine these variants. This analysis verified that 40 of these 49 CNVs (37 duplications, 3 deletions) were clearly segregating within at least one clonal cluster (Supplementary Figure 3). For example, one clonal population of seven closely related strains, collected together in 1966 from grape must in Sicily, have an average pairwise difference of only 19 SNPs (diversity π = 1.5×10^−6^). Notably, this collection showed four non-overlapping segregating duplications (Fig. 3c). This striking finding suggests that CNVs can arise or disappear frequently during evolution.

To examine whether this transience is a general feature of CNVs in this population, we quantified the variation in copy number of each CNV relative to mutations in the adjacent region of the genome. If a CNV was subject only to the same processes as these adjacent regions, we would expect a strong correlation between the total mutation in these regions and the total variation in copy number of the CNV. However, the variation in copy number of CNVs across the dataset was only weakly correlated with SNP variation in nearby regions of the genome (Spearman rank correlation ρ = 0.22, P = 0.041), indicating that CNVs are subject to additional or different evolutionary processes (Figure 2a). Furthermore, some CNVs showed high rates of variation within closely-related clusters relative to their variation in the rest of the dataset (Figure 2b and 2c, **Supplementary Table 2**, Supplementary Figure 4). Finally, we found that many CNVs represented the rare allele within the cluster, consistent with events that have short half-lives (Supplementary Figure 5). Taken together, these results indicate that CNVs are transient and variable features of the genome, even within extremely closely related strains.

**Figure 2.**
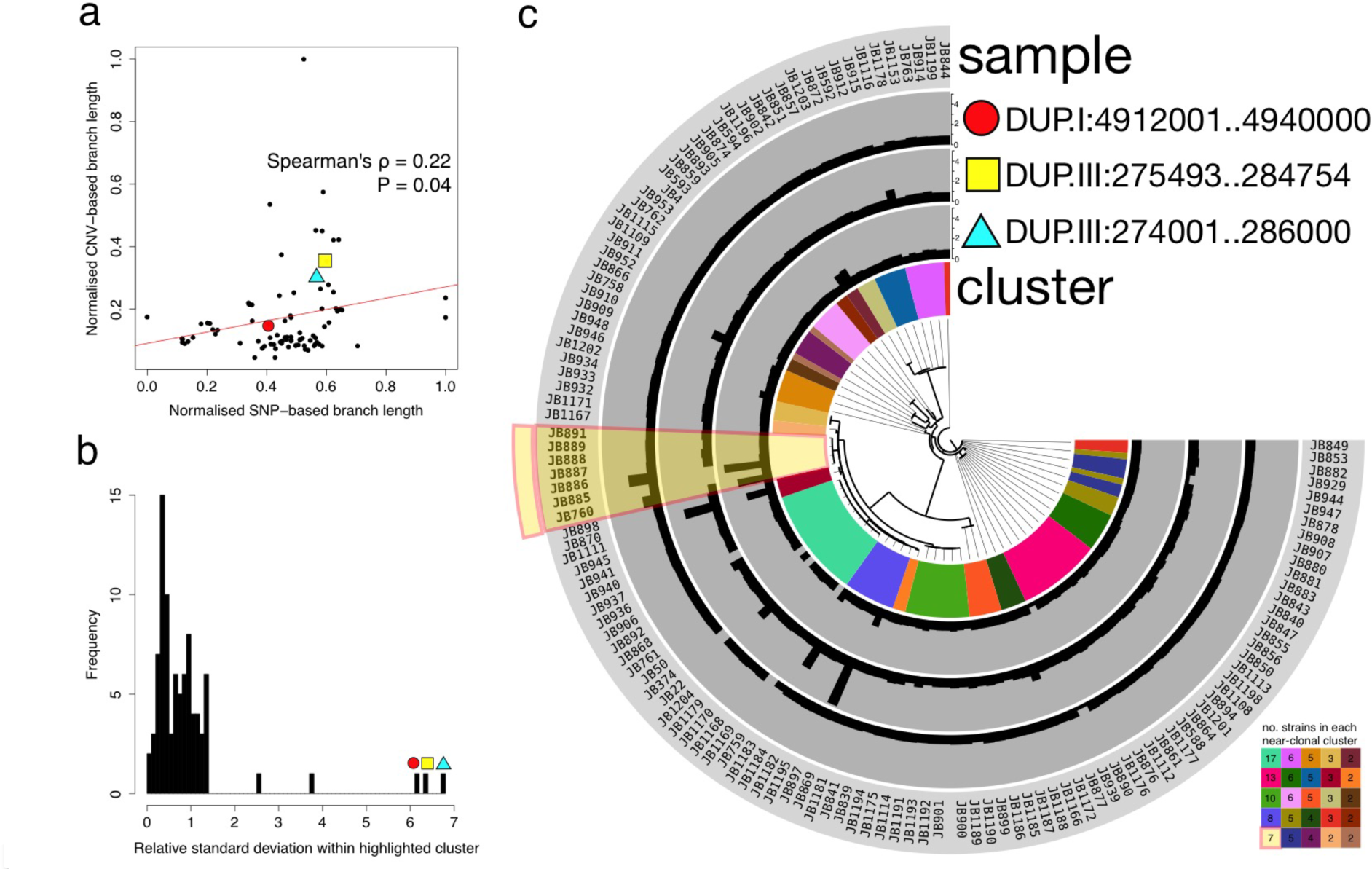
Copy number variants are transient within fission yeast. **(a)** For each of the 87 CNVs we calculated the genetic distance between strains using SNPs in the region around the CNV (20 kb up- and down-stream of the CNV, merged) as the total branch length from an approximate maximum-likelihood tree (*x*-axis, SNP-based branch length normalised to maximum value). We further calculated a CNV-based distance using the total branch length from a neighbour-joining tree constructed from Euclidean distances between strains based on their copy numbers (*y*-axis, CNV-based branch length normalised to maximum value). The weak correlation indicates that CNVs are subject to additional or different evolutionary processes. **(b)** Histogram of the standard deviation of each CNV within a near-clonal cluster (see also Figure 2a), relative to its standard deviation across strains not in the near-clonal cluster. Standard deviation is highly correlated with CNV-based branch length (Spearman rank correlation ρ = 0.90, P < 0.001) (Supplementary Figure 4b). The highlighted CNVs have unusually high rates of variation within this cluster compared to other clusters. **(c)** Copy number variation of these highlighted CNVs plotted on a SNP-based phylogeny (20kb up- and down-stream of the DUP.III:274001..286000 CNV) shows their relative transience within the cluster, as well as their variation across other near-clonal clusters. SNP-based phylogenies for the other two selected CNVs also do not separate the strains with different copy numbers (individual plots for each CNV across clusters for its corresponding SNP-based phylogeny are available as Supplementary data).

### Transient duplications affect gene expression

Partial aneuploidies of 500-700 kb in the *S. pombe* reference strain are known to alter gene expression levels within and, to some extent, outside of the duplicated region^31^. The naturally occurring duplications described here are typically smaller (median length: 21 kb), including an average of 6.5 genes. To examine whether naturally occurring CNVs have similar effects on gene expression, we examined eight pairs of closely related strains (<150 SNPs among each pair) that contained at least one unshared duplication (Figure 3, **Supplementary Table 3)**. Several of these strain pairs have been isolated from the same substrate at the same time, and all pairs are estimated to have diverged approximately 50 to 65 years ago (**Supplementary Table 3**). We assayed transcript expression from log phase cultures using DNA microarrays, each time comparing a duplicated to a non-duplicated strain from within the same clonal population. In seven of the eight strain pairs, the expression levels of genes within duplications were significantly induced, although the degree of expression changes between genes was variable (Figure 3c, Supplementary Figure 6). The increased transcript levels correlated with the increased genomic copy numbers, so that higher copy numbers produced correspondingly more transcripts (Spearman rank correlation ρ = 0.71, P = 0.014, Supplementary Figure 7). No changes in gene expression were evident immediately adjacent to the duplications (Supplementary Figure 7), suggesting that the local chromatin state was not strongly altered by the CNVs. This result not only confirms the previous observation that CNVs alter the gene expression levels, but more importantly it reveals large copy number differences between two genomes that are only 19 SNPs apart.

**Figure 3.**
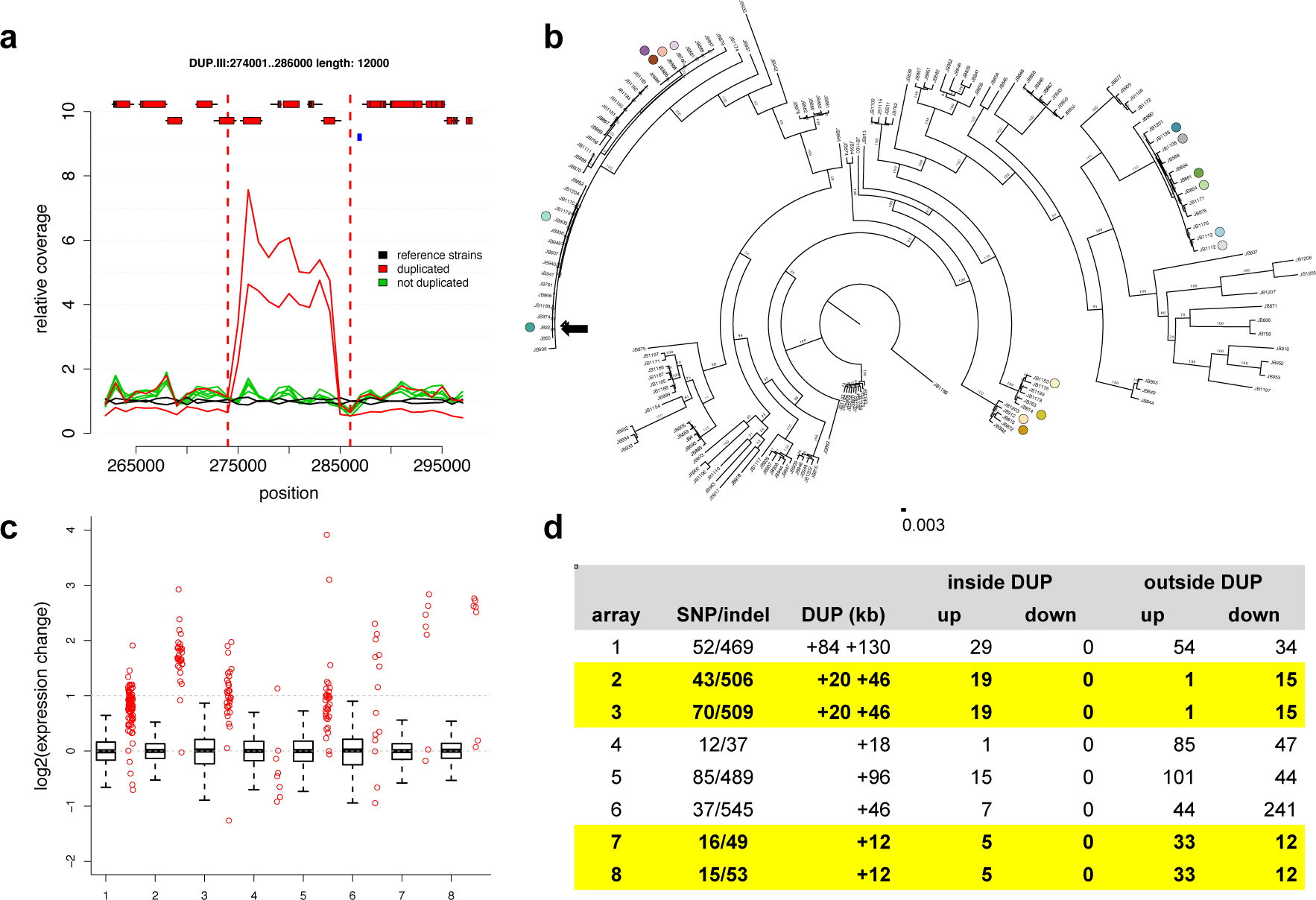
Transient duplications affect gene expression. (**a**) Duplications occur within near-clonal strains. Plot showing average read coverage in 1 kb windows for two clonal strains (JB760, JB886) with the duplication (red), five strains without duplication (green), and two reference strains (h^+^, and h-) (black). Genes (with exons as red rectangles) and retrotransposon LTRs (blue rectangles) are shown on top (see **Supplementary Table 3** for details). (**b**) Eight pairs of closely related strains, differing by one or more large duplications, selected for expression analysis. The tree indicates the relatedness of these strain pairs (dots colored as in d). The position of the reference strain (Leupold’s 972, JB22) is indicated with a black arrow. The scale bar shows the length of 0.003 insertions per site. (**c**) Gene expression increases for most genes within duplicated regions. For each tested strain pair, we show the relative gene expression (strains with duplication/strains without duplication) for all genes outside the duplication (as boxplot) and for all genes within the duplication (red strip chart). In all but one case (array 4), the genes within the duplication tend to be more highly expressed than the genes outside of the duplication (all Wilcoxon rank sum test P-values <1.5×10^−3^). (**d**) Summary of expression arrays 1-8, with strains indicated as colored dots (as in b), showing number of single-nucleotide polymorphism differences between strains (SNP), sizes of duplications in kb (DUP, where ‘+X +Y’ indicates two duplications with lengths X and Y, respectively). We show total numbers of induced (up) and repressed (down) genes, both inside and outside the duplicated regions. Arrays 2,3 and 7,8 (in yellow shading) are replicates within the same clonal population that contain the same duplications, so we list the number of up- and down-regulated genes that are consistent between both arrays. See **Supplementary Tables 3 and 4** for details.

Interestingly, some genes outside the duplicated regions also showed altered expression levels (Figure 3d, **Supplementary Table 4**). For example, two strain pairs differ by a single 12 kb duplication. Here, five of seven genes within the duplication showed induced expression, while 45 genes outside the duplicated region also showed consistently altered expression levels (38 protein-coding genes, 7 non-coding RNAs) (Figure 3d, arrays 7 and 8). As environmental growth conditions were tightly controlled, these changes in gene expression could be due to either compensatory effects of the initial perturbation caused by the 12kb duplication or changes that arise due to SNPs or indels that segregate between the strains (Supplementary Figure 6). We conclude that these evolutionary unstable duplications reproducibly affect the expression of distinct sets of genes and thus have the potential to influence cellular function and phenotypes.

### Copy number variants contribute to quantitative trait variance

To test whether SVs affect phenotypes, we examined the contributions of SNPs, CNVs and rearrangements to 227 quantitative traits (**Supplementary Table 5**), including 20 cell shape parameters, colony size on solid media assaying 42 stress and nutrient conditions^17^, 126 growth parameters in liquid media conditions^7^ and three biochemical parameters from wine ermentation^32^. For each phenotype, we used mixed model analysis to estimate the total proportion of variance explained by the additive contribution of genomic variants (the narrow-sense heritability).

When we determined heritability using only SNP data, estimates varied between 0% and 74% (median 30%). After adding CNVs and rearrangements to SNPs in a composite model, the estimated overall heritability increased for nearly all traits, explaining up to ~40% of trait variance (Figure 4a). This finding indicates that the CNVs and rearrangements can explain a substantial proportion of the trait variance. Using this composite model, we quantified the individual contributions of heritability best explained by SNPs, CNVs and rearrangements (Figure 4b). On average, SNPs explained 24% of trait variance, CNVs 7%, and rearrangements 4% (**Supplementary Table 5**). Analysis of simulated data confirmed that the contribution of CNVs could not be explained by linkage to causal SNPs alone (Supplementary Figure 6).

**Figure 4.**
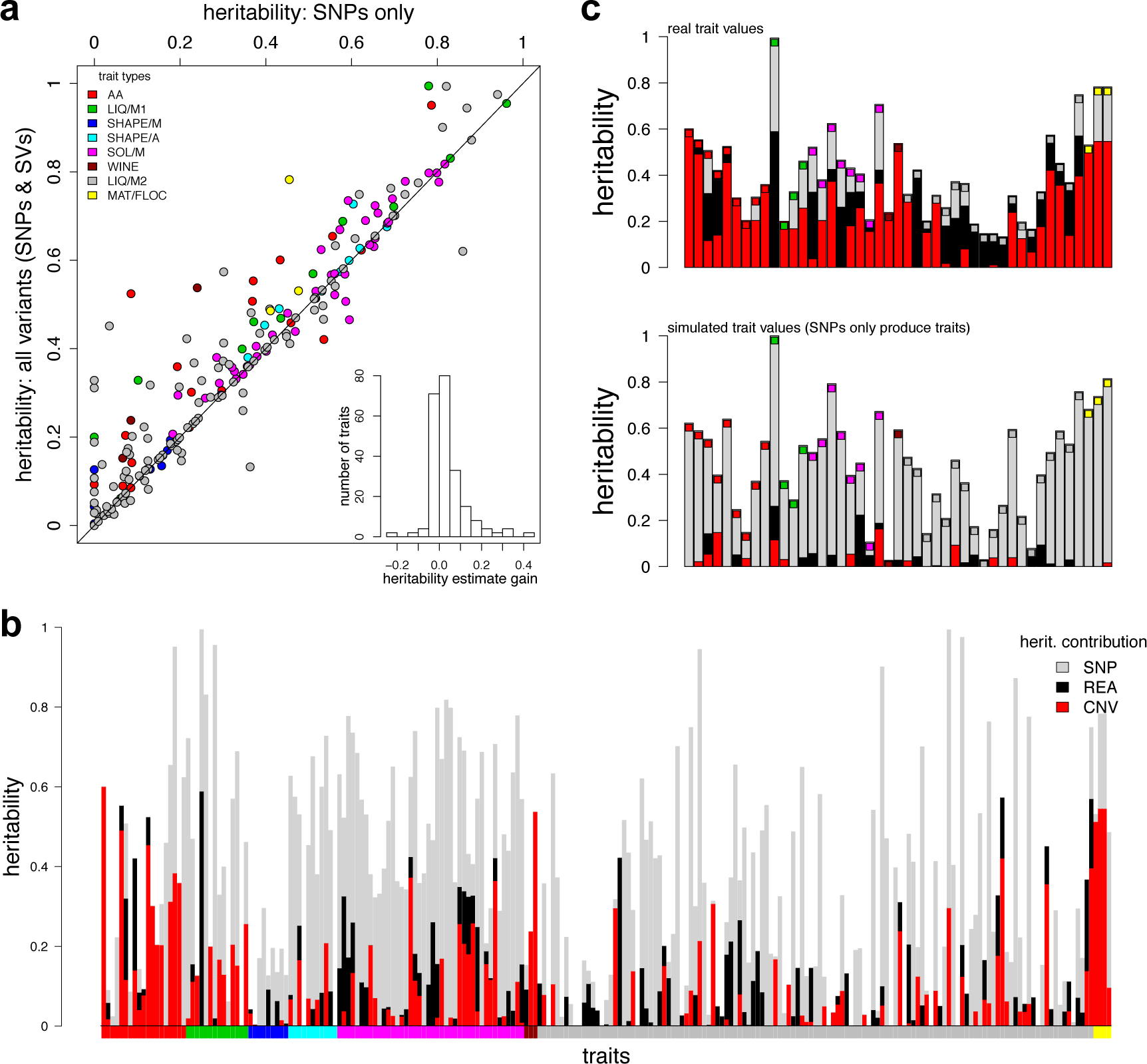
SVs contribute to quantitative traits. (**a**) Heritability estimates are improved by the addition of SVs. Heritability estimates for 227 traits (**Supplementary Table 5**), using only SNP data (x axis) range from 0 to 96% (median 29%). Adding SV calls (y axis) increases the estimates (median 34%), with estimates for some traits being improved up to a gain of 43% (histogram inset). The diagonal line shows where estimates after adding SVs are the same as those without (x=y). Inset: the distribution of the ‘gain’ in heritability after adding SV calls (median 0.4%, maximum 43%). Points are colored by trait types, according to legend top left. (**b**) The contributions of SNPs (grey), CNVs (red) and rearrangements (black) to heritability varied considerably between traits. Coloured bars along the *x* axis indicate the trait types. heritability estimates are in **Supplementary Table 5**. The panel below bars indicates trait types as in the legend for part (a). (**c**) Top panel, for some traits, SVs explained more of the trait variation than SNPs. Boxes are colored as legend in part (a). Lower panel, analysis of simulated data generated with assumption that only SNPs cause traits indicates that the contribution of SVs to trait variance is unlikely to be due to linkage.

Many trait measures gathered using the same method (e.g., growth on solid media, cell shape) are strongly correlated^17^. Thus, some groups of traits have consistently larger contributions from SVs (Figure 4b) than from SNPs alone. These traits include intracellular amino acid concentrations, growth under stress and several traits measured during wine fermentation (Figure 4c). Since many of these strains have been collected from fermentations (**Supplementary Table 6**), the substantial influence of CNVs may represent recent strong selection and adaptation to fermentation conditions that has occurred via recent CNV acquisition.

Our analysis of heritability showed that SNPs are generally able to capture most, but not all, of the genetic contribution of SVs (Figure 4). To examine whether trait-influencing SVs would be effectively detected from SNPs alone in this population, we examined the linkage of all 113 SVs with SNPs. We found that only 63 of these SVs (55%) are in strong linkage to SNPs (*r*^2^ >0.6), leaving 45% of the SVs weakly linked. This lack of linkage is consistent with SVs being transient, rather than persisting within haplotypes. Such weakly linked SVs may be missed in SNP-only association studies.

To examine this possibility, and to locate specific SVs that affect these traits, we performed mixed model genome-wide association studies, using all 68 SVs with minor allele counts >5 (i.e. occurring in at least 5 strains) as well as 139,396 SNPs and 22,058 indels with minor allele counts >5. Trait-specific significance thresholds for 5% familywise error rates were computed via permutation analysis, and were approximately 10^−4^ (SVs) and 10^−6^ (SNPs and indels). Nineteen SVs (28%) were significantly associated with traits (15 duplications, 5 deletions, 1 translocation), as well as 228 SNPs (0.16%), and 93 indels (0.42%) (**Supplementary Table 7**). SVs were associated with 20 different traits, including amino acid concentrations, mating traits, and stress resistance in solid and liquid media. Nine of these SVs were not strongly linked to SNPs (*r*^2^ < 0.6). The median effect size of these SVs was 14% (range 6-33%). While more detailed analyses of these associations will be required to confirm any particular association, our findings are consistent with the heritability analysis.

Collectively, these analyses indicate that even a small collection of SVs, most notably CNVs, can contribute substantially to quantitative traits. Thus, GWAS analyses conducted without genotyping SVs could fail to capture these important genetic factors.

### Structural variations contribute to intrinsic reproductive isolation

Crosses between *S. pombe* strains produce between <1% and 90% viable offspring^6,18^ We have previously shown that spore viability correlates inversely with the number of SNPs between the parental strains^17^. This intrinsic reproductive isolation may be due to the accumulation of Dobzhansky-Muller incompatibilities (variants that are neutral in one population, but incompatible when combined)^33,34^. However, genetically distant strains also accumulate SVs, which are known to lower hybrid viability and drive reproductive isolation^9^. In *S. pombe,* engineered inversions and translocations reduce spore viability by ~40%^6^. At present the impact of naturally occurring rearrangements, sequence divergence, and incompatible alleles in speciation within budding yeast is unclear^12–14,35,36^

To analyse intrinsic reproductive isolation in our population based on naturally occurring SVs, we examined the relationship between viability, SNPs and SVs. Both SV-distance (number of unshared SVs between parents) and SNP-distance inversely correlated with hybrid viability (Kendall correlation coefficients, SVs: **τ** = −0.26, P = 5.6 × 10^−3^, SNPs: **τ** = −0.35, P = 1.6×10^−4^) (Supplementary Figure 7). While inversions and translocations are known to lower hybrid viability as they affect chromosome pairing and segregation during meiosis^6,18,37^, CNVs are not expected to influence spore viability. Consistent with this view, there was no significant correlation between CNVs and viability (rearrangements, τ = −0.36, P = 2.0×10^−4^; CNVs, τ = −0.10, P = 0.28).

As the numbers of SNP and rearrangement differences between mating parents are themselves correlated (τ = 0.53, P = 1.3×10^−8^), we also estimated the influence of each factor alone using partial correlations. When either SNPs or rearrangements were controlled for, both remained significantly correlated with offspring viability (P = 0.04, P = 0.02, respectively) (Figure 5). Taken together, these analyses indicate that both rearrangements and SNPs contribute to reproductive isolation, but CNVs do not.

**Figure 5.**
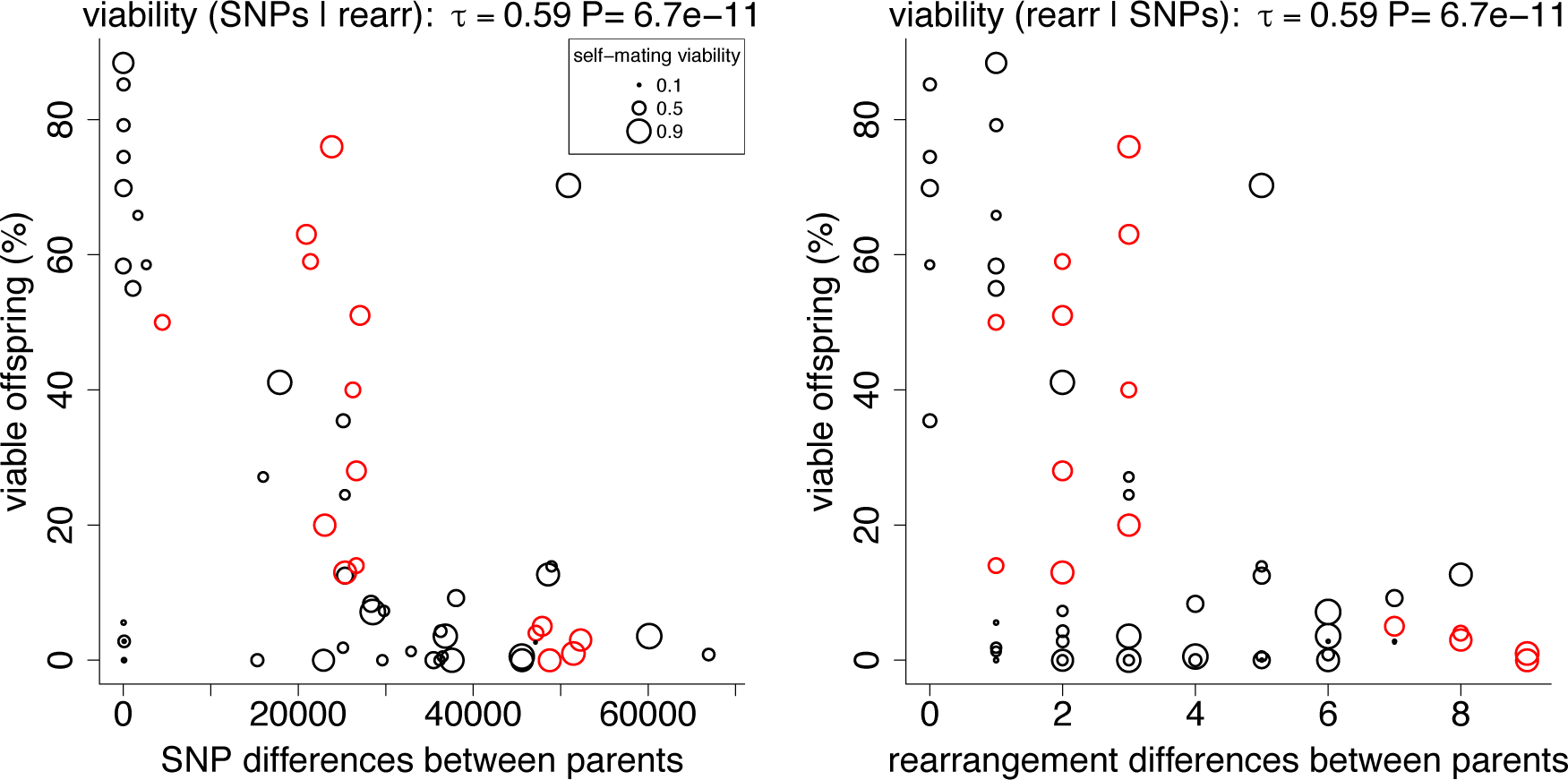
Both SNPs and rearrangements contribute to intrinsic reproductive isolation. Spore viability was measured from 58 different crosses from Jeffares, et al. ^17^ (black) or Avelar, et al. ^6^ (red), with each circle in the plots representing one cross. An additive linear model incorporating both SNP and rearrangement differences showed highly significant correlations with viability (P = 1.2×10^−6^, r^2^ = 0.39). Both genetic distances measured using SNPs and rearrangements (inversions and translocations) significantly correlated with viability when controlling for the other factor (Kendall partial rank order correlations with viability SNPs|rearrangements **τ** = −0.19, P = 0.038; rearrangements|SNPs **τ** = −0.22, P = 0.016). Some strains produce low viability spores even when self-mated with their own genotype. The lowest self-mating viability of each strain pair is indicated by circle size (see legend, smaller circles indicate lower self-mating viability) to illustrate that low-viability outliers tend to include such cases (see **Supplementary Table 8** for details).

## DISCUSSION

Here we present the first genome- and population-wide catalog of SVs among *S. pombe* strains. To account for the high discrepancy of available methods^25^, we applied a consensus approach to identify SVs (SURVIVOR), followed by rigorous filtering and manual inspection of all calls. We focused on high specificity (the correctness of the inferred SV) rather than high sensitivity (attempting to detect all SVs).

Our previous analyses of these strains, conducted without SV data^17^, attributed both trait variations and reproductive isolation to SNPs and/or small indels. Here we show that the small number of SVs we describe make substantial contributions to both of these factors. We demonstrate that CNVs (duplications and deletions) contribute significantly to our ability to describe quantitative traits, whereas variants that rearrange the order of the genome (inversions and translocations) produce much weaker effects on traits. In contrast, CNVs have no detectable influence on reproductive isolation, while rearrangements contribute substantially to reproductive isolation, similar to other species^10,38^

We show that CNVs and, to a lesser extent, rearrangements can produce substantial contributions to trait variation. These CNVs subtly alter the expression of genes within and beyond the duplications, and contribute considerably to quantitative traits. Within small populations, CNVs may produce larger effects on traits in the short term than SNPs, since their effect sizes can be substantial (SVs significant in GWAS have a mean effect size of 16% in this study). Within budding yeast, clearly measured effects of alterations to gene order in the DAL metabolic cluster^39^ and the lethality of some engineered rearrangements^40^ indicates that rearrangements can also effect phenotypic changes. Given the evidence for extensive ploidy and aneuploidy variation with budding yeasts, including clinical and industrial budding yeasts^29,41,42^, SVs can be expected to have considerable impacts on phenotypic variation these fungi.

In this context, it is striking that CNVs appear to be transient within the clonal populations that we studied. Our analysis is consistent with experimental studies with budding yeast, indicating that both rearrangements and CNVs may be gained or lost at rates in excess of point mutations. For example, frequent gain of duplications has been observed in laboratory cultures of *S. pombe*, where spontaneous duplications suppress *cdc2* mutants at least 100 times more frequently than point mutations. These suppressor strains lose their duplications with equal frequency^43^, indicating reversion of alleles. Similarly, duplications frequently occur during experimental evolution with budding yeast. This instability is likely facilitated by repeated elements, which are unstable within both budding and fission yeast genomes^45–48^, which is also supported by the enrichment of SVs in our population near retrotransposon LTRs (Supplementary Figure 8). Though we do not examine the stability of rearrangements, there is also evidence for their instability. Transposon-mediated rearrangements are highly dynamic in laboratory cultures during selection^49^,^50^, and show elevated mutation rates at subtelomeric regions^51^.

This analysis also has relevance for human diseases, since *de novo* CNV formation in the human genome occurs at a rate of approximately one CNV/10 generations^52^, and CNVs are known to contribute to a wide variety of diseases^4^. Indeed, both the population genetics and the effects of SVs within *S. pombe* seem similar to human, in that CNVs are associated with stoichiometric changes on gene expression, and SVs are in weak linkage with SNPs^53,54^, and therefore may be badly tagged by SNPs in GWAS studies. We show that CNVs and rearrangements in fission yeast not only rapidly emerge, but substantially contribute to quantitative traits independent of weakly linked SNPs. These findings highlight the need to identify SVs when describing traits using GWAS, and indicate that a failure to call SVs can lead to an overestimation of the impact of SNPs to traits or contribute to the problem that large proportions of the heritable component of trait variation are not discovered in GWAS (the ‘missing heritability’). We observed a clear example of this effect in two winemaking traits, where heritability was entirely due to SVs.

In summary, we show that different types of SVs are transient within populations of fission yeast, where they alter gene expression, impact phenotypes and can lead to reproductive isolation.

## METHODS

### Performance assessment of SV callers using simulated data

To identify filtering parameters for DELLY, LUMPY and Pindel for the *S. pombe* genome, we simulated seven datasets (s1-s7) of 40x coverage with a range of different SV types and sizes (**Supplemental Table 7**). The simulated read sets contained sequencing errors (0.4%), SNPs and indels (0.1%) within the range of actual data from *S. pombe* strains and between 30 and 170 SVs. These data sets were produced by modifying the reference genome using our in-house software (SURVIVOR, described below), and simulating reads from this genome with Mason software^55^.

After mapping the reads and calling SVs, we evaluated the calls. We defined a SV correctly predicted if: i) the simulated and reported SV were of the same type (e.g. duplication), ii) were predicted to be on same chromosome, and iii) their start and stop locations were with 1 kb. We then defined caller-specific thresholds to optimize the sensitivity and false discovery rate (FDR) for each caller. FDRs on the simulated data were low: DELLY (average 0.13), LUMPY (average 0.06) and Pindel (average 0.04).

Selecting calls that were present in at least two callers further reduced the FDR (average of 0.01). DELLY had the highest sensitivity (average 0.75), followed by SURVIVOR (average 0.70), LUMPY (average 0.62) and Pindel (0.55). We further used simulated data to assess the sensitivity and FDR of our predictions. cn.mops was evaluated with a 2 kb distance for start and stop coordinates. Our cn.mops parameters were designed to identify large (above 12 kb) events and thus did not identify any SVs simulated for s1-s6. Details of simulations and caller efficacy are provided in **Supplementary Table 9**.

### SURVIVOR (StructURal Variant majorIty VOte) Software Tool

We developed the SURVIVOR tool kit for assessing SVs for short read data that contains several modules. The first module simulates SVs given a reference genome file (fasta) and the number and size ranges for each SV (insertions, deletions, duplications, inversions and translocations). After reading in the reference genome, SURVIVOR randomly selects the locations and size of SV following the provided parameters. Subsequently, SURVIVOR alters the reference genome accordingly and prints the so altered genome. In addition, SURVIVOR provides an extended bed file to report the locations of the simulated SVs.

The second module evaluates SV calls based on a variant call format (VCF) file ^56^ and any known list of SVs. A SV was identified as correct if i) they were of same type (e.g. deletion); ii) they were reported on same chromosome, and iii) the start and stop coordinates of the simulated and identified SV were within 1 kb (user definable).

The third module of SURVIVOR was used to filter and combine the calls from three VCF files. In our case, these files were the results of DELLY, LUMPY and Pindel. This module includes methods to convert the method-specific output formats to a VCF format. SVs were filtered out if they were unique to one of the three VCF files. Two SVs were defined as overlapping if they occur on the same chromosome, their start and stop coordinates were within 1 kb, and they were of the same type. In the end, SURVIVOR produced one VCF file containing the so filtered calls. SURVIVOR is available at github.com/fritzsedlazeck/SURVIVOR.

### Read mapping and detection of structural variants

Illumina paired-end sequencing data for 161 *S. pombe* strains were collected as described in Jeffares, et al. ^17^, with the addition of Leupold’s reference 975 *h*^+^ (JB32) and excluding JB374 (known to be a gene-knockout version of the reference strain, see below). Leupold’s 968 *h*^*90*^ and Leupold’s 972 *h*^−^ were included as JB50 and JB22, respectively (**Supplementary Table 6**). For all strains, reads were mapped using NextGenMap (version 0.4.12)^57^ with the following parameter (-X 1000000) to the *S. pombe* reference genome (version ASM294v2.22). Reads with 20 base pairs or more clipped were extracted using the script *split_unmapped_to_fasta.pl* included in the LUMPY package (version 0.2.9)^25^ and were then mapped using YAHA (version 0.1.83)^58^ to generate split-read alignments. The two mapped files were merged using Picard-tools (version 1.105) (http://broadinstitute.github.io/picard), and all strains were then down-sampled to 40× coverage using Samtools (version 0.1.18)^59^.

Subsequently, DELLY (version 0.5.9, parameters: “ -q 20 -r”)^26^, LUMPY (version 0.2.9, recommended parameter settings)^25^ and Pindel (version 0.2.5a8, default parameter)^27^ were used to independently identify SVs in the 161 strains using our SURVIVOR software. This included merging any variants of the same type (duplication, deletion *etc*) whose start and end coordinates where within 1 kb. Merging was justified by the finding that most allele calls were close to the defined call (only 5% of start or end positions were >300nt from the defined consensus boundary). We then retained all variants predicted by at least two methods. These SVs calls were genotyped using DELLY.

To identify further CNVs, we ran cn.MOPS^24^ with parameters tuned to collect large duplications/deletions as follows: read counts were collected from bam alignment files (as above) with *getReadCountsFromBAM* and WL=2000, and CNVs predicted using *haplocn.mops* with minWidth= 6, all other parameters as default. Hence, the minimum variant size detected was 12 kb. CNV were predicted for each strain independently by comparing the alternative strain to the two reference strains (JB22, JB32) and four reference-like strains that differed from the reference by less than 200 SNPs (JB1179, JB1168, JB937, JB936).

After CNV calling, allele calling was achieved by comparing counts of coverage in 100bp windows for the two reference strains (JB22, JB32) to each alternate strain using custom R scripts. Alleles were called as non-reference duplications if the one-sided Wilcoxon rank sum test p-values for both JB22 and JB32 vs alternate strain were less than 1×10^-10^ (showing a difference in coverage) and the ratio of alternate/reference coverage (for both JB22 and JB32) was >1.8 (duplications), or <0.2 (deletions). Manual inspection of coverage plots showed that the vast majority of the allele calls were in accordance with what we discerned by eye. These R scripts were also used to examine CNVs predicted to be segregating within clusters (clonal populations). All such CNVs were examined in all clusters that contained at least one non-reference allele call (**Supplementary Table 10**).

Finally, we manually mapped two large duplications that did not satisfy these criteria (DUP.I:2950001..3190000, 240kb and DUP.I:5050001..5560000, 510kb — both singletons in JB1207), but were clearly visible in chromosome-scale read coverage plots (Supplementary Figure 9).

### Reduction of false discovery rate

This filtering produced 315 variant calls. However, because 31 of these 315 (~10%) were called within the two reference strains (JB22, JB32), we expected that this set still contained false positives. To further reduce the false positive rate, we looked for parameters that would reduce calls made in reference strains (JB22 and JB32) but not reduce calls in strains more distantly related to the reference (JB1177, JB916 and JB894 that have 68223, 60087 and 67860 SNP differences to reference^17^). The reasoning was that we expected to locate few variants in the reference, and more variants in the more distantly related strains. This analysis showed that paired end support, repeats and mapping quality were of primary value.

We therefore discarded all SVs that had a paired end support of 10 or less. In addition, we ignored SVs that appeared in low mapping quality regions (i.e. regions where reads with MQ=0 map) or those where both start and end coordinates overlapped with previously identified retrotransposon LTRs^17^.

Finally, to ensure a high specificity call set, these filtered SVs were manually curated using IGV ^60^ (**Supplementary Tables 11,12**). We assigned each SVs a score (0: not reliable, 1: unclear, 2: reliable based on inspection of alignments through IGV). We utilized different visualizations from IGV to identify regions were pairs of the reads mapped to different loci, for example, which we interpreted as possible artefacts. Overall, we investigated whether the alignments of the breakpoints and reads in close proximity had a reliable mapping in terms of mapping quality and clearness of the distortions of the pairs. Only calls passing this manual curation as reliable (score 2) were included in the final data set of 113 variants utilized for all further analyses. These filtering and manual curation steps reduced our variant calls substantially, from 315 to 113. At this stage only 1/113 (~1%) of these variants was called within the two standard reference strains (Leupolds’s *h+* and *h-,* JB22 and JB32 in our collection).

### PCR validation

PCR analysis was performed to confirm 10 of the 11 inversions and all 15 translocations from the curated data set. One inversion was too small to examine by PCR (INV.AB325691:6644..6784, 140 nt). Primers were designed using Primer3^61^ to amplify both the reference and alternate alleles. PCR was carried out with each primer set using a selection of strains that our genotype calls predict to include at least one alternate allele and at least one reference allele (usually 6 strains). Products were scored according to product size and presence/absence (**Supplementary Tables 13,14**).

Inversions: 9/10 variants were at least partially verified by either reference or alternate allele PCR (3 variants were verified by both reference and alternate PCRs), and 7/10 inversions also received support from BLAST (see below). Translocations: 10/15 were at least partially verified by either reference or alternate allele PCR (5/15 variants were verified by both reference and alternate PCRs). One additional translocation received support from BLAST (see below), meaning that 11/15 translocations were supported by PCR and/or BLAST. Three of the four translocations that could not be verified were probably nuclear copies of mitochondrial genes (NUMTs)^62^, because one breakpoint was mapped to the mitochondrial genome. Details of the 113 curated variants are presented in Supplementary Table 15.

### Validation by BLAST of *de novo* assemblies

We further assessed the quality of the predicted breakpoints for the inversions and translocations by comparing them to the previously created *de novo* assemblies for each of the 161 strains^17^. To this end, we created blast databases for the scaffolds of each strain that were >1kb. We then created the predicted sequence for 1 kb around each junction of the validated 10 inversions and 15 translocations. These sequences were used to search the blast databases using BLAST+ with --gapopen 1 --gapextend 1 parameters. We accepted any blast hsp with a length >800 bp as supporting the junction (because these must contain at least 300 bp at each side of the break point). Four inversions and three translocations gained support from these searches (Supplementary File Tables2-PCR.xlsx).

### Knockout strain control

Our sample of sequenced strains included one strain (JB374) that is known to contain deletions of the *his3* and *ura4* genes. Our variant calling and validation methods identified only two variants in this strain, both deletions that corresponded to the positions of these genes, as below:

*his3* gene location is chromosome II, 1489773-1488036, deletion detected at II:1488228-1489646.

*ura4* gene location is chromosome III, 115589-116726, deletion detected at III:115342-117145.

This strain was not included in the further analyses of the SVs.

### Microarray expression analysis

Cells were grown in YES (Formedium, UK) and harvested at OD_600_ =0,5. RNA was isolated followed by cDNA labeling^63^. Agilent 8 × 15K custom-made *S. pombe* expression microarrays were used. Hybridization, normalization and subsequent washes were performed according to the manufacturer’s protocols. The obtained data were scanned and extracted using GenePix and processed for quality control and normalization using in-house developed R scripts. Subsequent analysis of normalized data was performed using R. Microarray data have been submitted to ArrayExpress (accession number E-MTAB-4019). Genes were considered as induced if their expression signal after normalization was >1.9, and repressed if <0.51.

### Time to most recent common ancestor (TMRCA) estimates

Previously, based on the genetic distances between these strains and the ‘dated tip’ dating method implemented in BEAST^64^, we have estimated the divergence times between all 161 *S. pombe* strains sequenced^17^. To determine the TMRCA for pairs of strains, we re-examined the BEAST outputs using FigTree to obtain the medium and 95% confidence intervals.

### SNP and indel calling

SNPs were called as described^17^. Insertions and deletions (indels) were called in 160 strains using stampy-mapped, indel-realigned bams as described previously^17^. We accepted indels that were called by both the Genome Analysis Toolkit HaplotypeCaller^65^ and Freebayes^66^, and then genotyped all these calls with Freebayes.

Briefly, indels were called on each strains bam with HaplotypeCaller, and filtered for call quality >30 and mapping quality >30 (bcftools filter --include 'QUAL>30 && MQ>30'). Separately, indels were called on each strains bam with Freebayes, and filtered for call quality >30. All Freebayes vcf files were merged, accepting only positions called by both Freebayes and HaplotypeCaller. These indels were then genotyped with Freebayes using a merged bam (containing reads from all strains), using the --variant-input flag for Freebayes to genotyped only the union calls. Finally indels were filtered for by score, mean reference mapping quality and mean alternate mapping quality >30 (bcftools filter --include 'QUAL>30 && MQM>30 & MQMR>30'). These methods identified 32,268 indels. Only 50 of these segregated between Leupold’s h^−^ reference (JB22) and Leupold’s h^90^ reference (JB50), whereas 12109 indels segregated between the JB22 reference and the divergent strain JB916.

### Heredity and GWAS

We selected 53 traits that contained at least values from 100 strains^17^, and so included multiple individuals from within clonal populations (growth rates on 42 different solid media and 11 cell shape characters measured with automated image analysis). Trait values were normalized using a rank-based transformation in R, for each trait vector *y*, normal.*y* =qnorm(rank(*y*)/(1+length(*y*))). Total heritability, and the contribution of SNPs, CNVs and rearrangements were estimated using LDAK (version 5.94)^67^, with kinship matrices derived from all SNPs, 146 CNVs, and 15 rearrangements. All genotypes, including CNVs were encoded as binary values (1 or 0) for heritability and GWAS. To assess whether the contribution of CNVs could be primarily due to linkage with causal SNPs, we simulated trait data using the --make-phenos function of LDAK with the relatedness matrix from all SNPs, assuming that all variants contributed to the trait (--num-causals −1). We made one simulated trait data set per trait, for each of the 53 traits, with total heritability defined as predicted from the real data. We then estimated the heritability using LDAK, including the joint matrix of SNPs, CNVs and rearrangements. To assess the extent to which the contribution of SNPs to heritability was overestimated, we performed another simulation using the relatedness matrix from the 87 segregating CNVs alone, and then estimated the contribution of SNPs, CNVs and rearrangements in this simulated data as above.

Genome-wide associations were performed with LDAK (version 5) using default parameters. To account for the unequal relatedness of strains, we used a kinship matrix derived from all 172,368 SNPs called previouslyJeffares, et al. ^17^. Association analysis was used to find associations between traits, testing SVs, SNPs and indels with a minor allele count ≥5. Analysis was run separately for 68 SVs, 139,396 SNPs and 22,058 indels (each used the kinship derived from all SNPs). We examined the same 53 traits as for the heritability analysis (above). For each trait, we carried out 1000 permutations of trait data, and define the 5^th^ percentile of these permutations as the trait-specific P-value threshold.

### Model details for Heritability and GWAS Analysis

To estimate the heritability contribution of SNPs, we computed a kinship matrix (K_SNP_) using all 172,368 SNPs that we had discovered in our previous published analysis^17^ (elements of this matrix represent pairwise allelic correlations across all SNPs)^67^, onto which we regressed the phenotypic values assuming the following model:

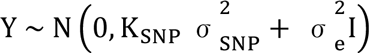

We estimated the two variance components, 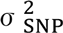 and 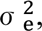 using REML (restricted maximum likelihood), based on which our estimates of the heritability of SNPs is

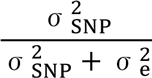

To estimate the heritability of CNVs and rearrangements, we repeated this analysis using instead K_CNV_ then K_REA_, computed using only 146 segregating CNVs and 15 segregating rearrangements, respectively.

We additionally considered the model

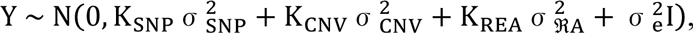

Having estimated the four variance components, again using REML, the relative contributions of SNPs, CNVs and rearrangements are, respectively,

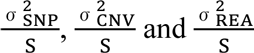

where 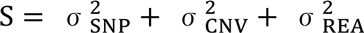

To test the specificity of this analysis, we generated phenotypes for which only one predictor type contributed (e.g., only SNPs), then analyzed using the individual and joint models above, which allowed us to assess how accurately we can distinguish between contributions of different predictor types.

For the mixed model association analysis, we used the same the SNP kinship matrix. As the predictors (variants that we examined for effects on a trait), we chose to analyse SNPs, indels and SVs with a minor allele count ≥5 (68 SVs, 139396 SNPs and 22,058 indels).

Then for each predictor X_j_ we considered the model

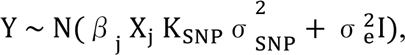

where *β* _j_ is the effect size of predictor X_j_

Having solved using REML, we used a likelihood ratio test (comparing to the null model (*β* _j_ = 0) to assess whether *β* _j_is significantly non-zero. Each of these analyses used the kinship derived from all SNPs.

### Offspring viability and genetic distance

Cross spore viability data and self-mating viability were collected from previous analyses^6,17^. The number of differences between each pair was calculated using vcftools vcf-subset ^56^, and correlations were estimated using R, with the ppcor package. When calculating the number of CNVs differences between strains, we altered our criteria for ‘different’ variants (to merge variants whose starts and ends where within 1 kb), and merged CNVs if their overlap was >50% and their allele calls were the same.

### Transience analysis

For each CNV, we extracted all SNPs from 20 kb upstream and 20 kb downstream. 86/87 CNVs showed variation in these regions (DUP.MT:1..19382 was the only CNV with no corresponding SNPs). We then used these concatenated SNPs to build a local SNP-based tree with FastTree (version 2.1.9)^68^. To build a CNV-based tree from the copy number variation in each CNV region, we used a neighbour-joining tree estimation based on the Euclidean distances between strains.

The total branch length of the CNV-based tree was strongly correlated (Spearman rank correlation ρ=0.90, P <0.001) with the standard deviation of copy number variation (Supplementary Figure 4). We therefore used this standard deviation to define a relative rate of transience for each cluster, σ_rc_ = σ_ic_/σ_oc_ where σ_ic_ and σ_oc_ are the within cluster and without cluster standard deviations respectively, meaning that CNVs which were highly relatively transient within a given cluster would have high values of σ_rc_. This was used to select the three CNVs visualised in Figure 2c. See Supplementary Table 2 for all values of σ_rc_, Supplementary Figure 4 for visualization as heatmap). Visualisations of all 86/87 CNVs with their SNP-based phylogenies are available at: https://figshare.com/projects/fission_yeast_structural_variation/15798.

Circle plots were used to visualize the variation in copy number over the SNP-based phylogeny for each CNV using Anvi’o (version 2.0.3)^69^.

## Supplementary data

SNP, indel and SVs calls, genotypes and copy numbers are available on figshare at: https://figshare.com/projects/fission_yeast_structural_variation/15798

Array data is available at ArrayExpress, accession number: E-MTAB-4019.

## Acknowledgments

We thank Günter Klambauer for advice on cn.MOPS and Michael C. Schatz for helpful discussions and comments on the manuscript. F.S. was supported through National Science Foundation awards (DBI-1350041) and National Institutes of Health award (R01-HG006677). D.J., M.H., C.R. were supported by a Wellcome Trust Senior Investigator Award to J.B. (grant 095598/Z/11/Z). J.B. was supported by a Royal Society Wolfson Research Merit Award.

## Author contributions

DJ, FS, CD and JB conceived and developed the study. DJ, LS, CJ and FS conducted the bioinformatics analysis. DJ designed the laboratory work. FS designed and implemented SURVIVOR. DS contributed to analysis of heritability and GWAS. CR and MH produced the expression array analysis. MH conducted PCR validation of variants. JB provided the bulk of funding for personnel and research costs. DJ, FS, CJ, LS, FB, CD and JB wrote the manuscript.

## Supplementary Figures

**Supplementary Figure 1.**
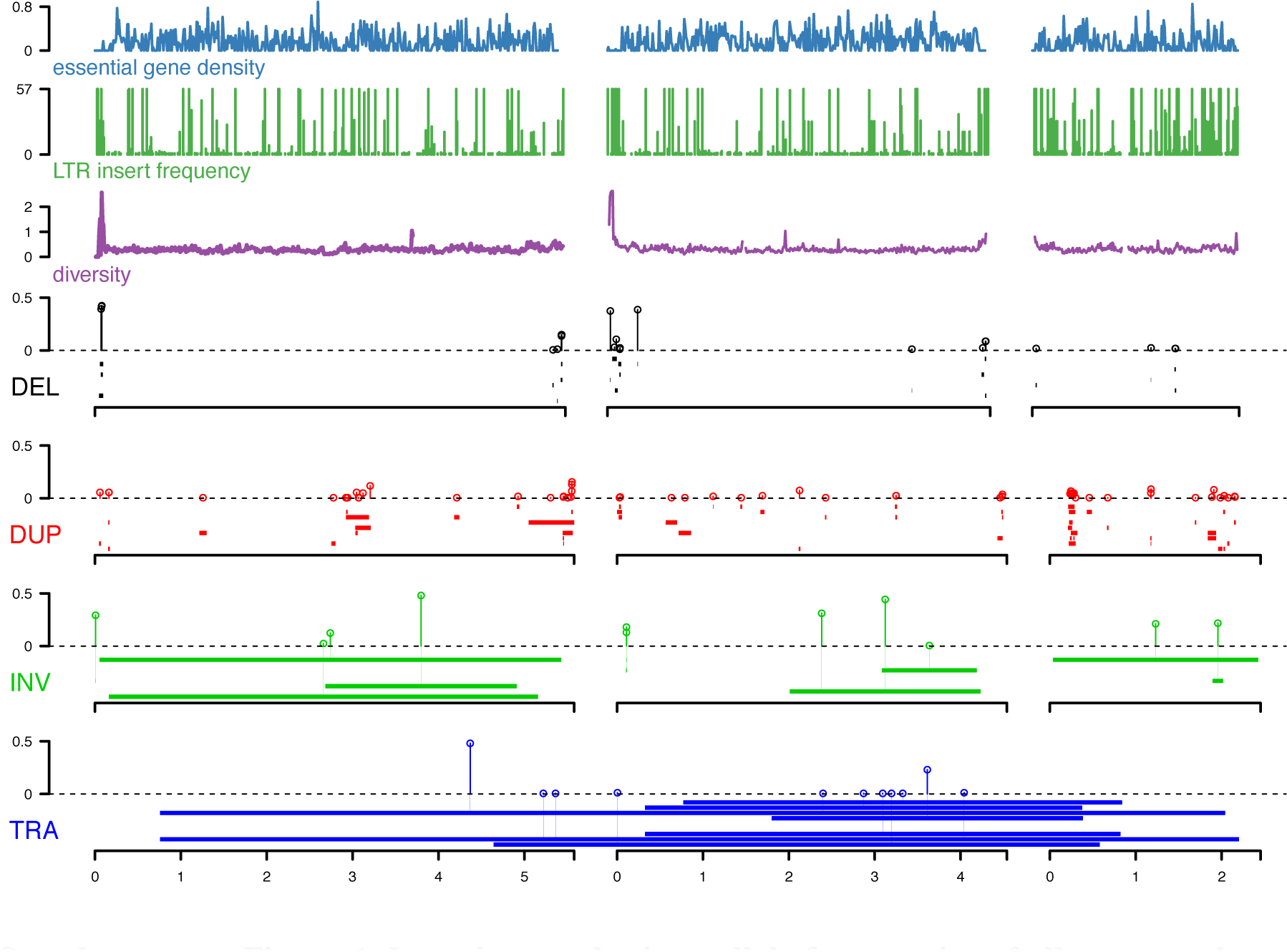
Locations and minor allele frequencies of all structural variants in curated data set. Each of the three chromosomes is indicated by black bar, with scale (in megabases) at bottom. From top (same data as Fig 1): density of essential genes (blue), locations of *Tf*-type retrotransposons (green), and diversity (π, average pairwise diversity from SNPs, purple). Bar heights for deletions and duplications are proportional to minor allele frequency, the scale for retrotransposons is the frequency of the insertion in the 57 non-clonal strains. Diversity and retrotransposon were calculated from 57 non-clonal strains as described in Jeffares, et al. ^17^. Below, we show different types of SVs: deletions (black), duplications (red), inversions (green) and translocations (blue). The vertical lines terminating with open circles above dotted lines emit from the mid-point of each SV and indicate the minor allele frequencies in the population of 161 strains.

**Supplementary Figure 2.**
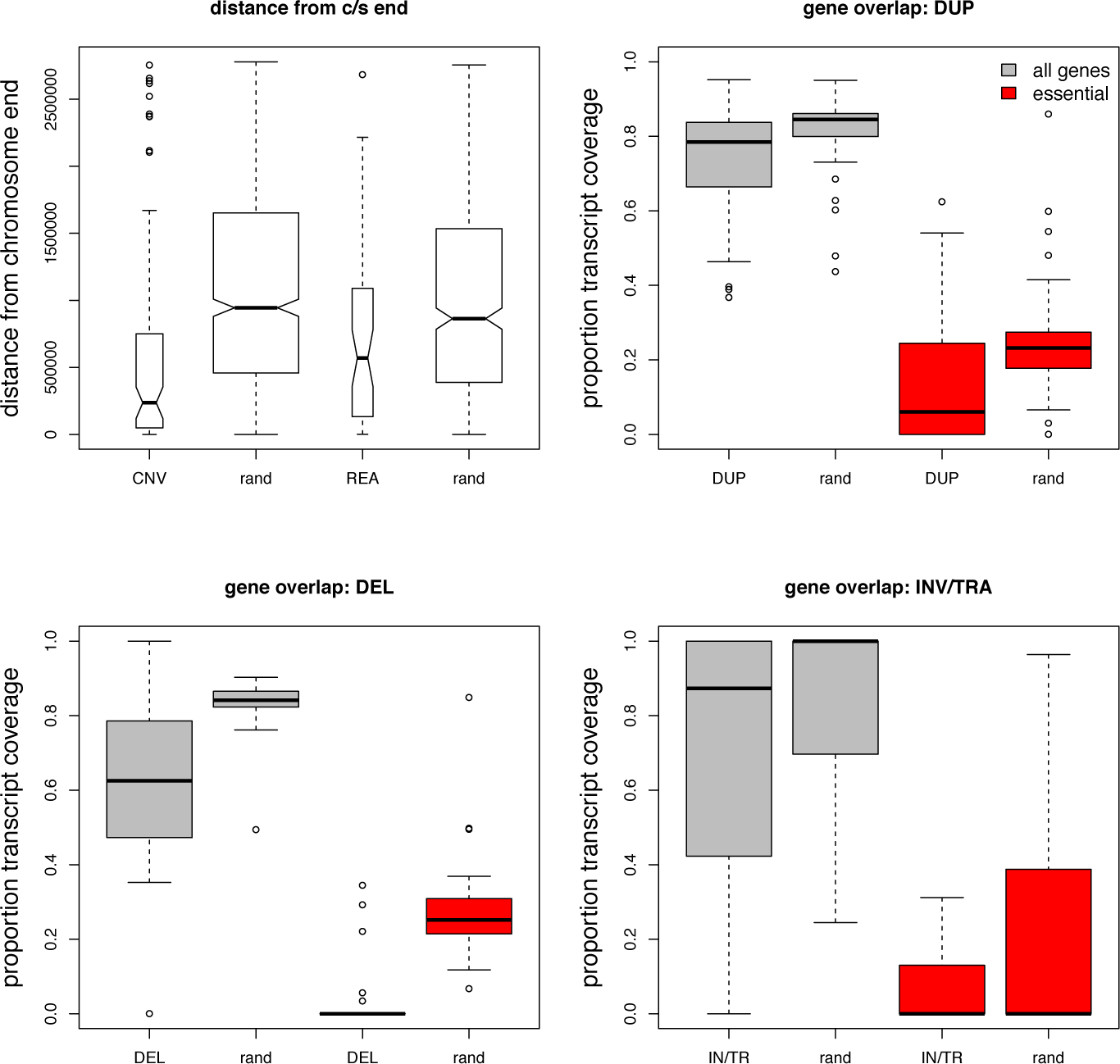
Structural variations are biased towards chromosome ends and to low gene density regions. Top left panel, both CNVs and rearrangements are biased towards the ends of chromosomes. CNVs; median distance to chromosome ends 236 kb compared to chromosome- and size-matched random sites 944 kb, Wilcoxon rank sum test P = 1.3 × 10^−11^, rearrangements median distance 569 kb *vs,* matched random 863 kb, Wilcoxon test P = 0.03). All other panels, calculated proportion of each duplication and deletion that contained all protein-coding or essential genes. Box plots show the distributions of these proportions for all genes (grey), and proportion of coverage by essential genes (red), compared to the null distribution (rand). All comparisons were significantly less than the null distributions (Wilcoxon rank sum test, P-values <1.6 × 10^−4^). The same analysis was performed with the junctions of inversions and translocations, by calculating the transcript coverage in the region 500 bp up- and down-stream of the predicted start and end junctions. These rearrangements are slightly biased away from genes (P = 1.9×10^−3^), but not significantly biased away from essential genes (P >0.05). The null distributions were determined by selecting 10 regions for each actual variant/junction that were the same size, and were placed in random positions on the same chromosome and calculating the gene coverage of these regions. Essential genes were those with the Fission Yeast Phenotype Ontology term defined as FYPO:0002061 (“inviable”) in PomBase.

**Supplementary Figure 3.**
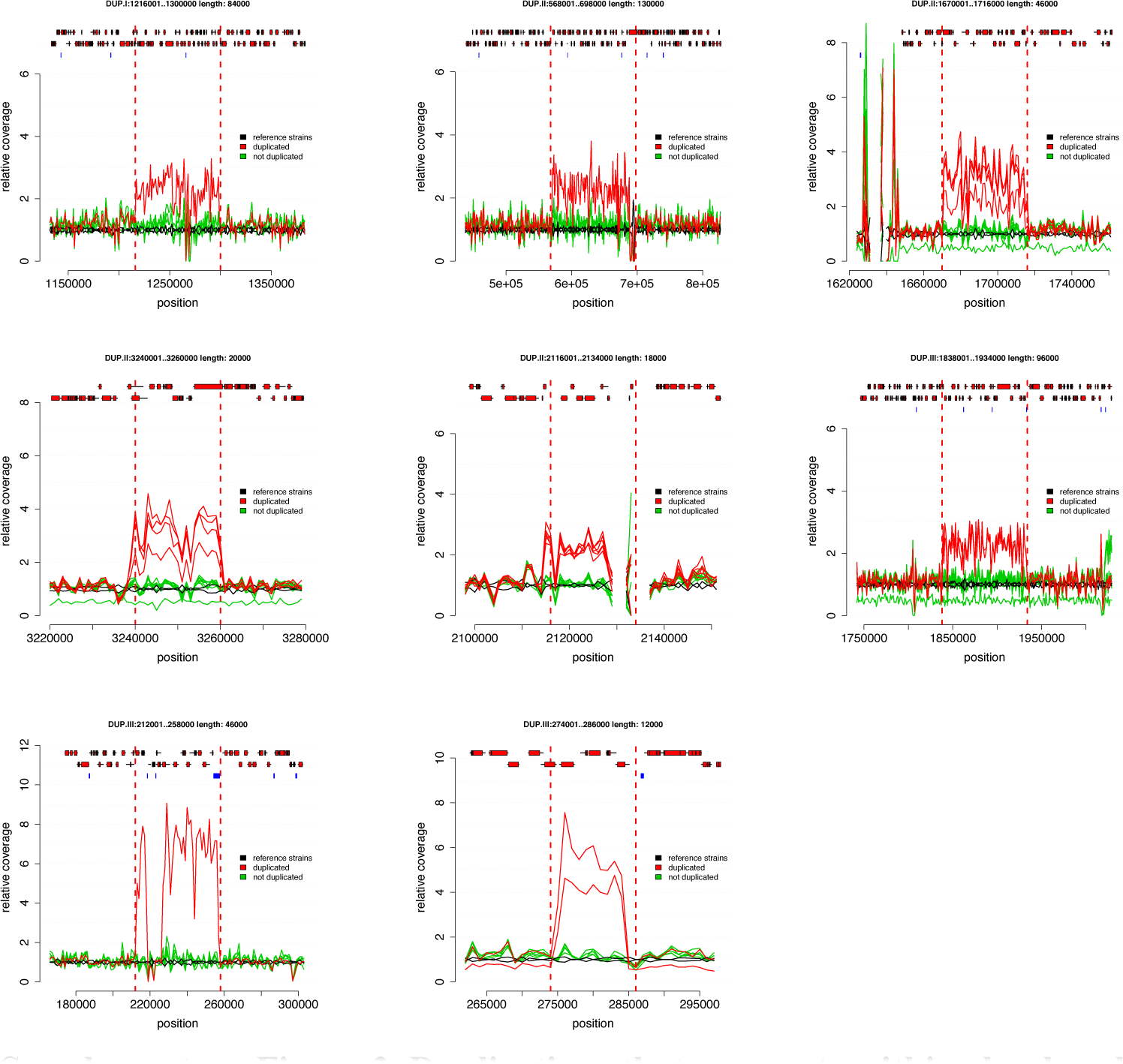
Duplications that segregate within closely related strains. Plots show the average coverage in 1 kb non-overlapping windows for strains with a duplication (red) and all closely related strains without duplication (green); all these strains differ by <150 SNPs. The coverage of the two standard reference strains (*h*^+^ and *h*^−^) is shown in black. Top row, from left: variant DUP.I:1216001..1300000 (cluster 12, from Japan in 57), DUP.II:568001..698000 (cluster 12), DUP.II:1670001..1716000 (cluster 2, unknown origin), second row DUP.II:3240001..3260000 (cluster 2), DUP.II:2116001..2134000 (cluster 1, includes reference strain from French grapes in 1947), DUP.III:1838001..1934000 (cluster 2, various locations 1921-22). Bottom row: DUP.III:212001..258000 (cluster 6, Jamaica/USA), and DUP.III:274001..286000 (cluster 5, Sicily 1966). Genes are shown on top of plots with exons as red rectangles and retrotransposon LTRs as blue rectangles.

**Supplementary Figure 4.**
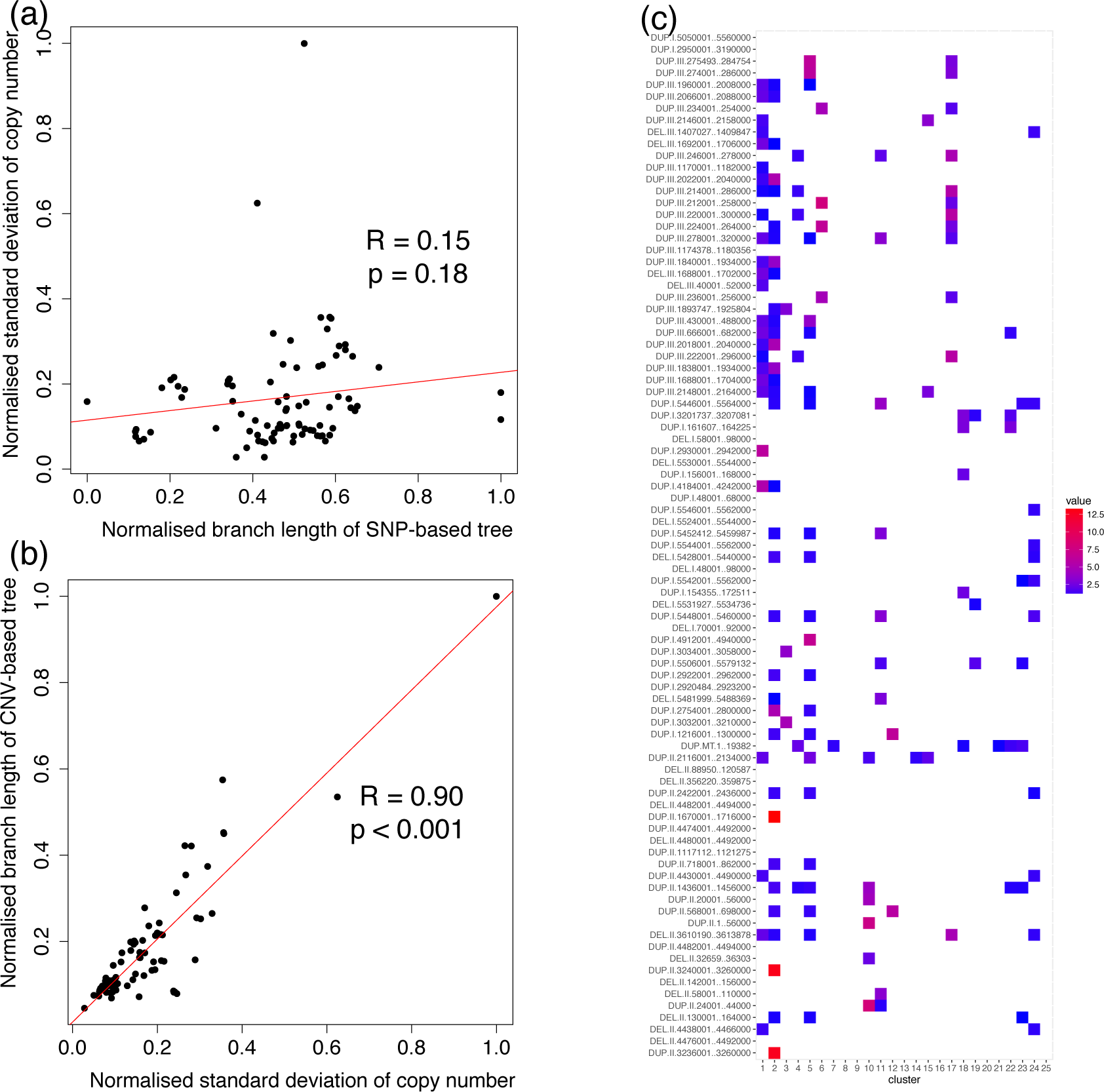
Relative standard deviation of copy number variation within clusters. (**a**) Standard deviation of copy number for a CNV across the dataset is only weakly correlated with the total branch length of SNP-based phylogeny from the 20kb up- and down-stream phylogeny. **(b)** Standard deviation is highly correlated with the branch length of a CNV-based neighbour-joining tree. **(c)** The relative standard deviation of each CNV within each identified cluster of strains (<150 SNPs apart) relative to its change in rest of the dataset. For clarity, all relative standard deviations <1 are shown as white.

**Supplementary Figure 5.**
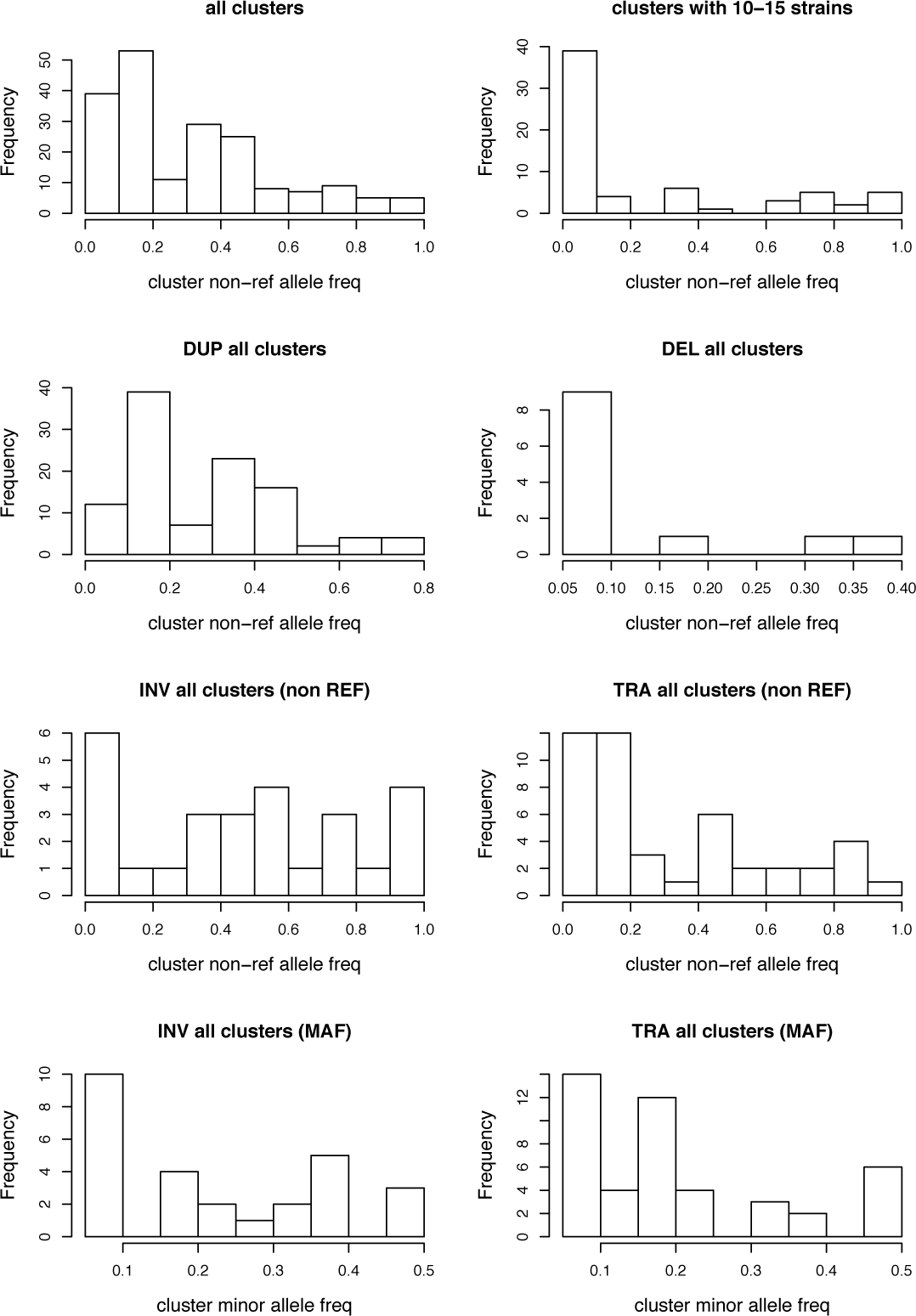
Copy number variants are usually rare alleles within clonal populations. Clonal clusters, or clonal populations all differ by < 150 SNPs. In rows, from top left; we show the within-cluster frequency of the non-reference allele for all SVs, which is skewed to rare alleles. Limiting this analysis to cluster with 10 to 15 strains highlights the low frequency of non-reference alleles. Second row; CNVs (duplications and deletions) are skewed to rare alleles, because the non-reference allele is usually the derived allele. Third row; inversions and translocations are not skewed to the non-reference allele, but here non-reference alleles are not necessarily the derived allele. Bottom row; the *minor allele* of inversions and translocations, however, is skewed to rare alleles.

**Supplementary Figure 6.**
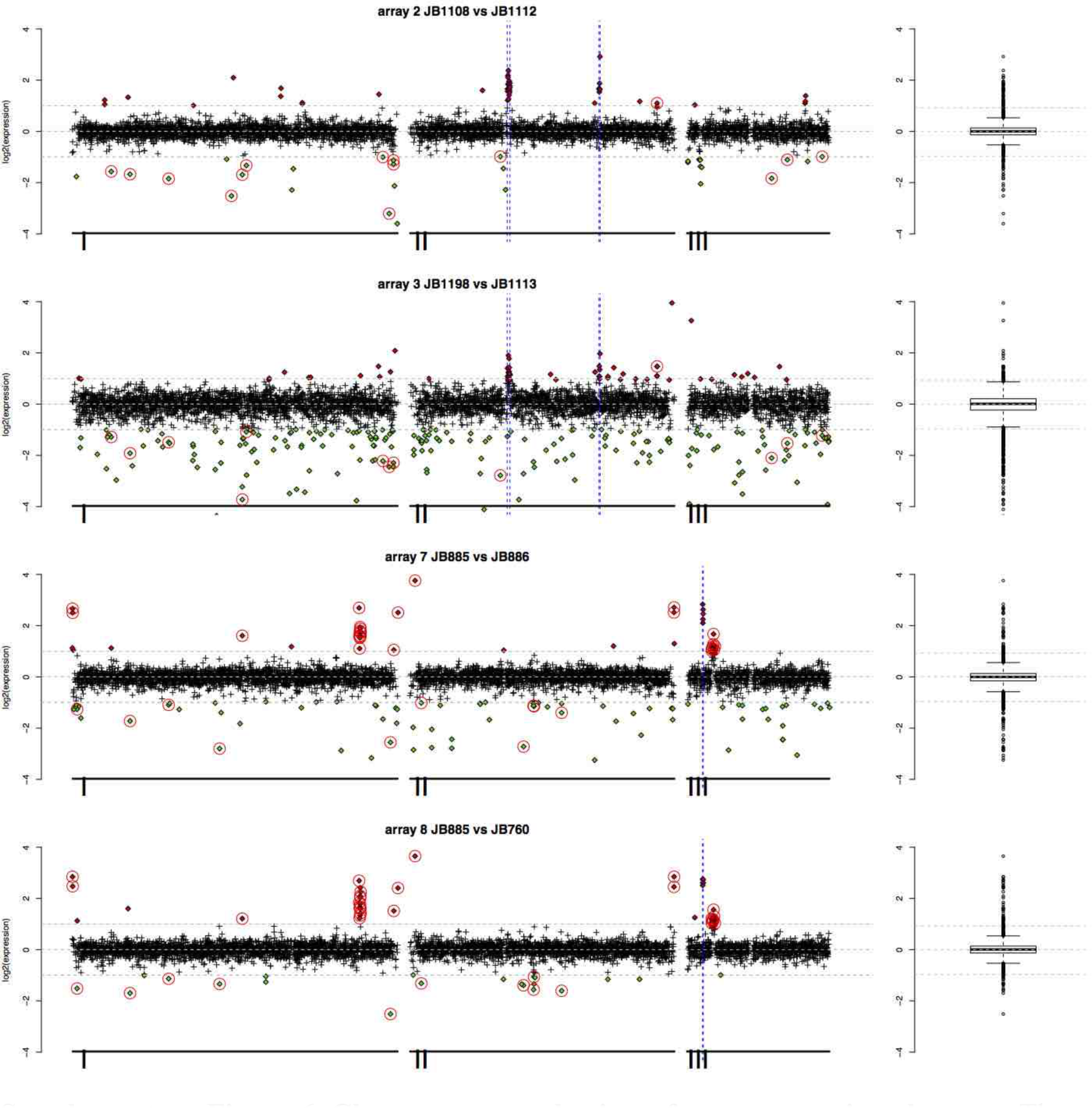
Chromosome-scale view of gene expression changes. The relative gene expression levels (strain1/strain2) for arrays 2 and 3, and arrays 7 and 8 are shown with their positions on the three chromosomes. Filled circles indicates genes that we consider to be up-regulated (red) or repressed (green). Those highlighted with open red circles are consistently altered in both arrays (either 2+3, or 7+8). The blue lines show where the segregating duplications are. Box plots at right show the spread of data.

**Supplementary Figure 7.**
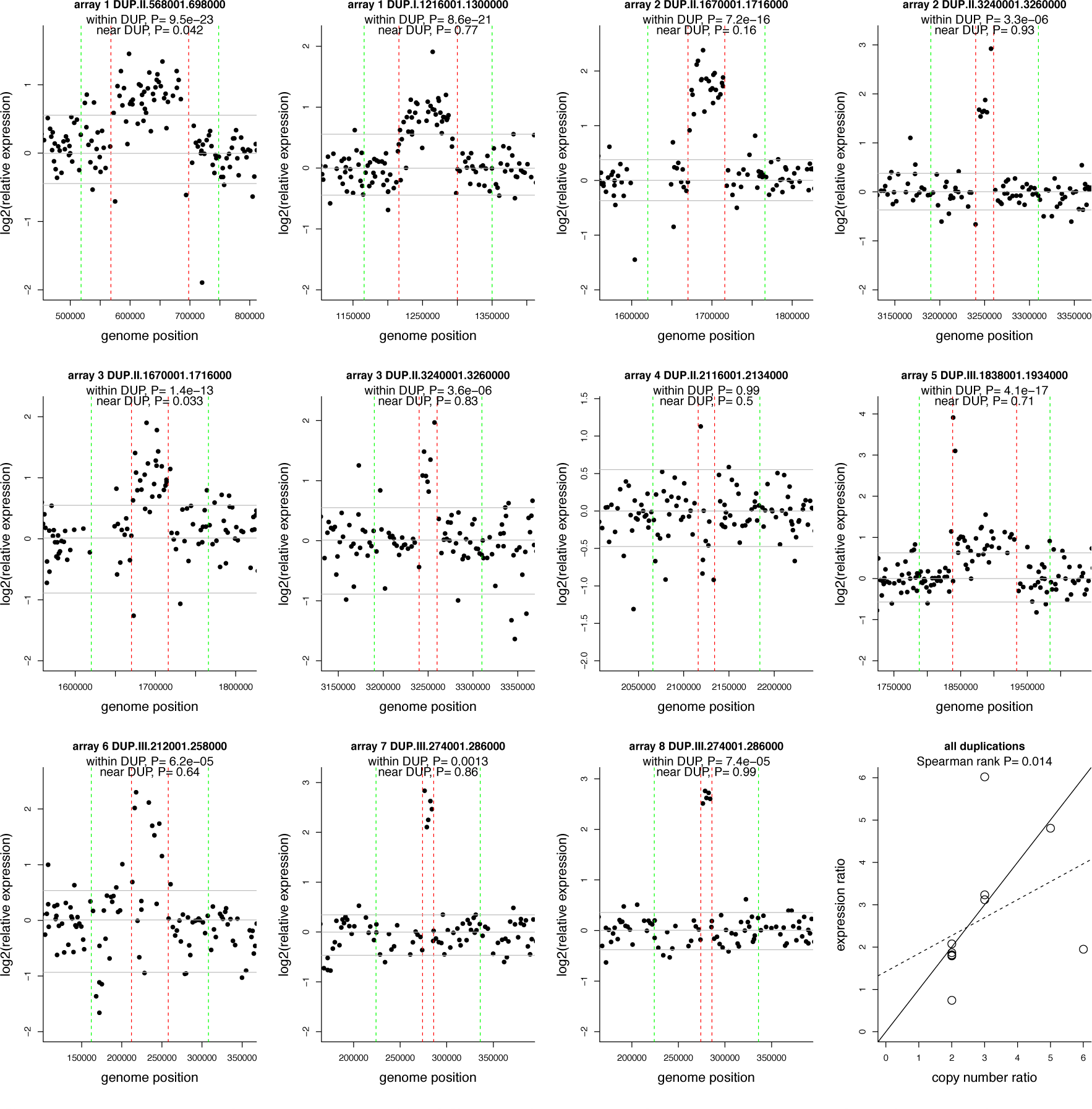
No significant increase in gene expression immediately adjacent to duplications. For each duplication examined with DNA arrays, we show the relative expression (strain 1 *vs* strain 2) near the duplication. P-values show the support for the genes within the duplication (red vertical lines), or the 50 kb adjacent to the duplication (green vertical lines) being more highly expressed than all other genes in the chromosome (one-sided Wilcoxon rank sum tests). The grey horizontal lines show the 5^th^, 50^th^ and 95^th^ percentiles for gene expression data on the chromosome. The bottom right panel shows that the median increase in expression level within a duplication correlates with the increase in genomic copy number. The solid back line shows the expected increase for the 1:1 correspondence between genomic copy number and relative expression (the line *y*=*x*), and the dashed line shows the linear model for the data. Copy number and relative expression change are correlated (Spearman rank correlation ρ = 0.71 and P = 0.014).

**Supplementary Figure 6.**
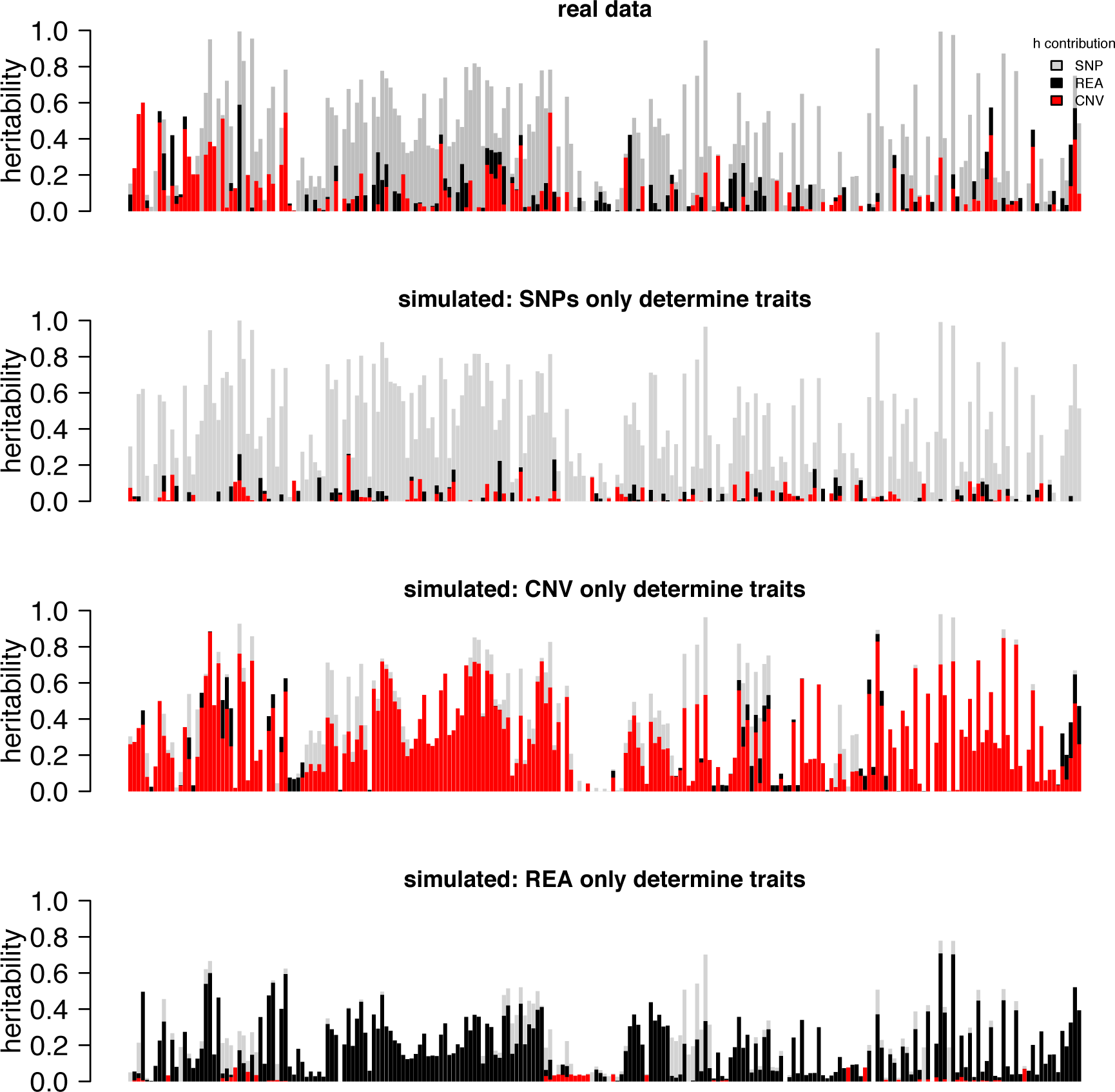
Contributions of SNPs, CNVs and rearrangements to traits. Top panel: for 227 traits, we show the total heritability estimated by the combination of 243,289 SNPs (green), 87 CNVs (red), and 26 rearrangements (grey). We then simulated data that was entirely due to the effects of SNPs (second panel), entirely due to the effects of CNVs (next panel) or entirely due to the effects of rearrangements (lower). In the second panel (entirely due to the effects of SNPs), the contribution of CNVs and rearrangements are artefacts, but these are relatively minor. This analysis indicates that the estimates are not strongly affected by linkage.

**Supplementary Figure 7.**
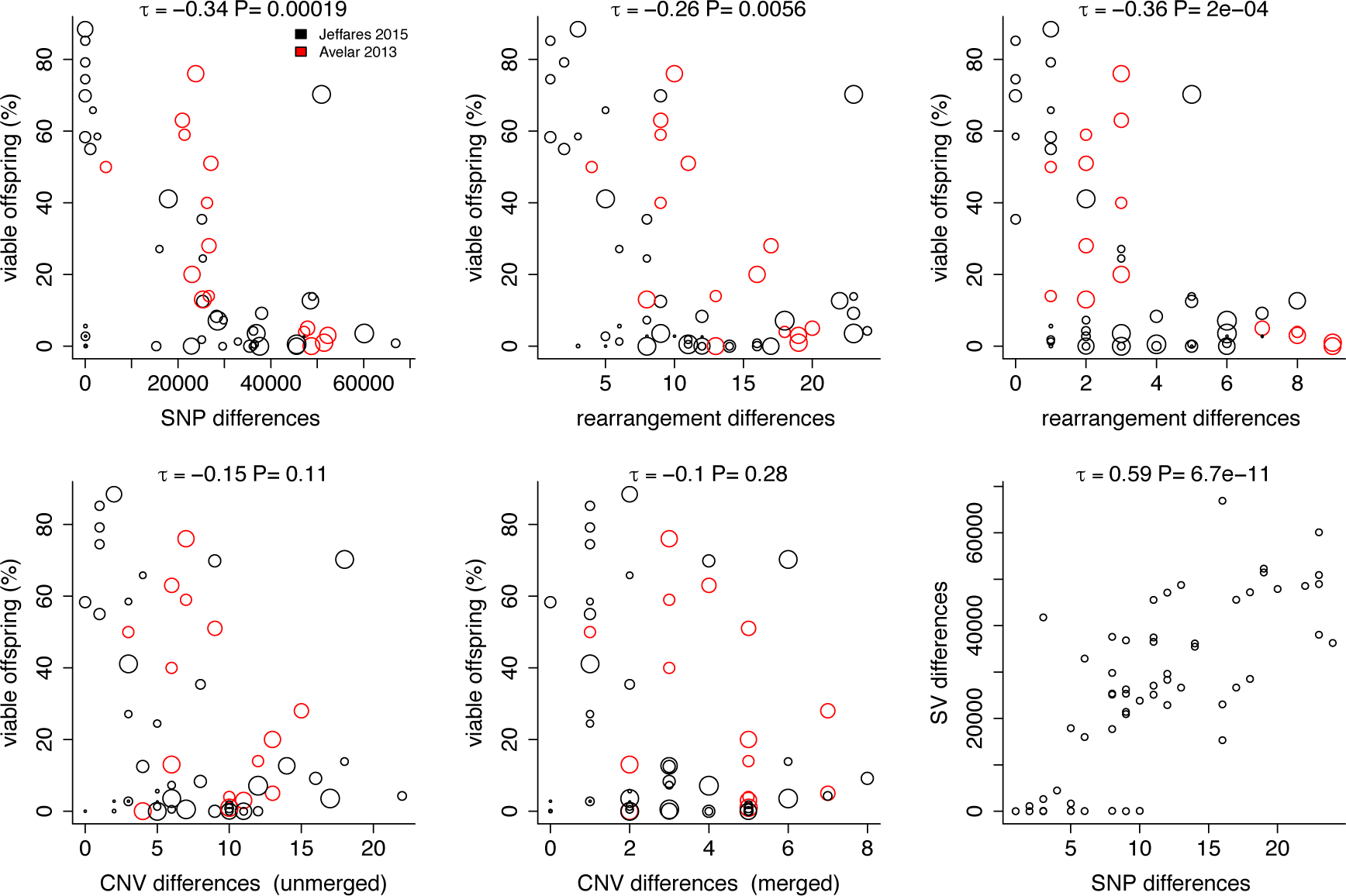
Correlations between spore viability, parental SNP-genetic distance and parental SV-genetic distance. Spore viability was measured for 58 crosses in total, including data from both Jeffares, et al. ^17^ (black) and Avelar, et al. ^6^ (red), with each circle representing one cross. Unmerged CNV differences count any CNV as being different between parents when either start or end coordinates are more than 1 kb apart. Because this definition can cause us to count largely overlapping events as ‘different’, we also counted ‘merged’ differences where two CNVs were considered different only if their overlap was >50% of the total of both variants. This approach will exclude nested CNVs. CNV-genetic distance is not significantly correlated with viability in either case.

**Supplementary Figure 8.**
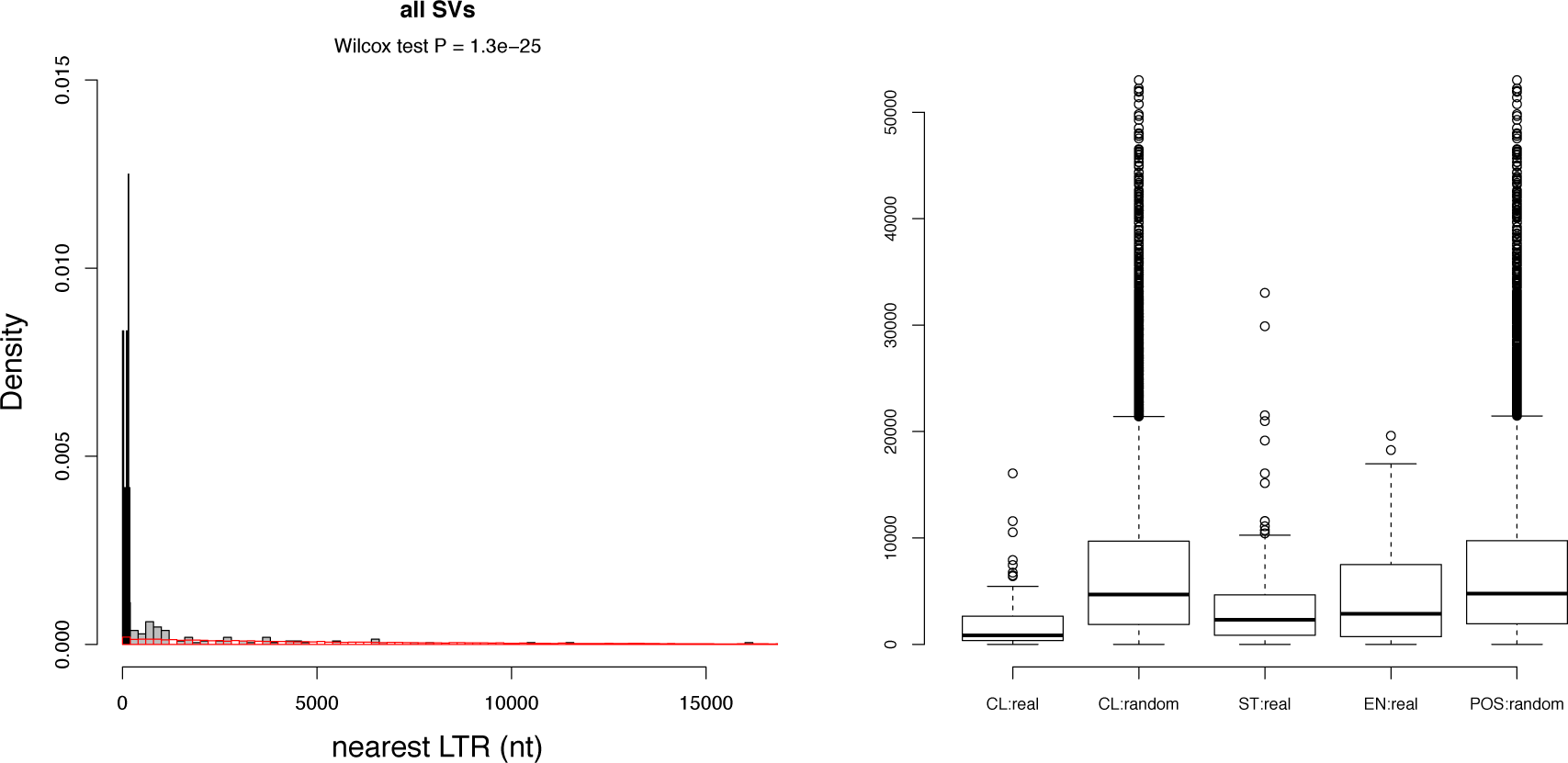
SVs are enriched close to retrotransposon LTRs. For all SVs, we computed the closest distance of start or end coordinates to any LTR discovered previously^17^. As a control, we compute the closest distance of 10 random coordinates on the same chromosome. Left: the distributions of distances for real SVs (grey), those that are within 200nt (black) or random coordinates (red). Right: using the same analysis, we show the closest distance of real SVs (CL: real), and random coordinates (C: random). We also show that both start and end coordinates of SVs (ST:real, EN:real) are closer than random positions (POS:random).

**Supplementary Figure 9.**
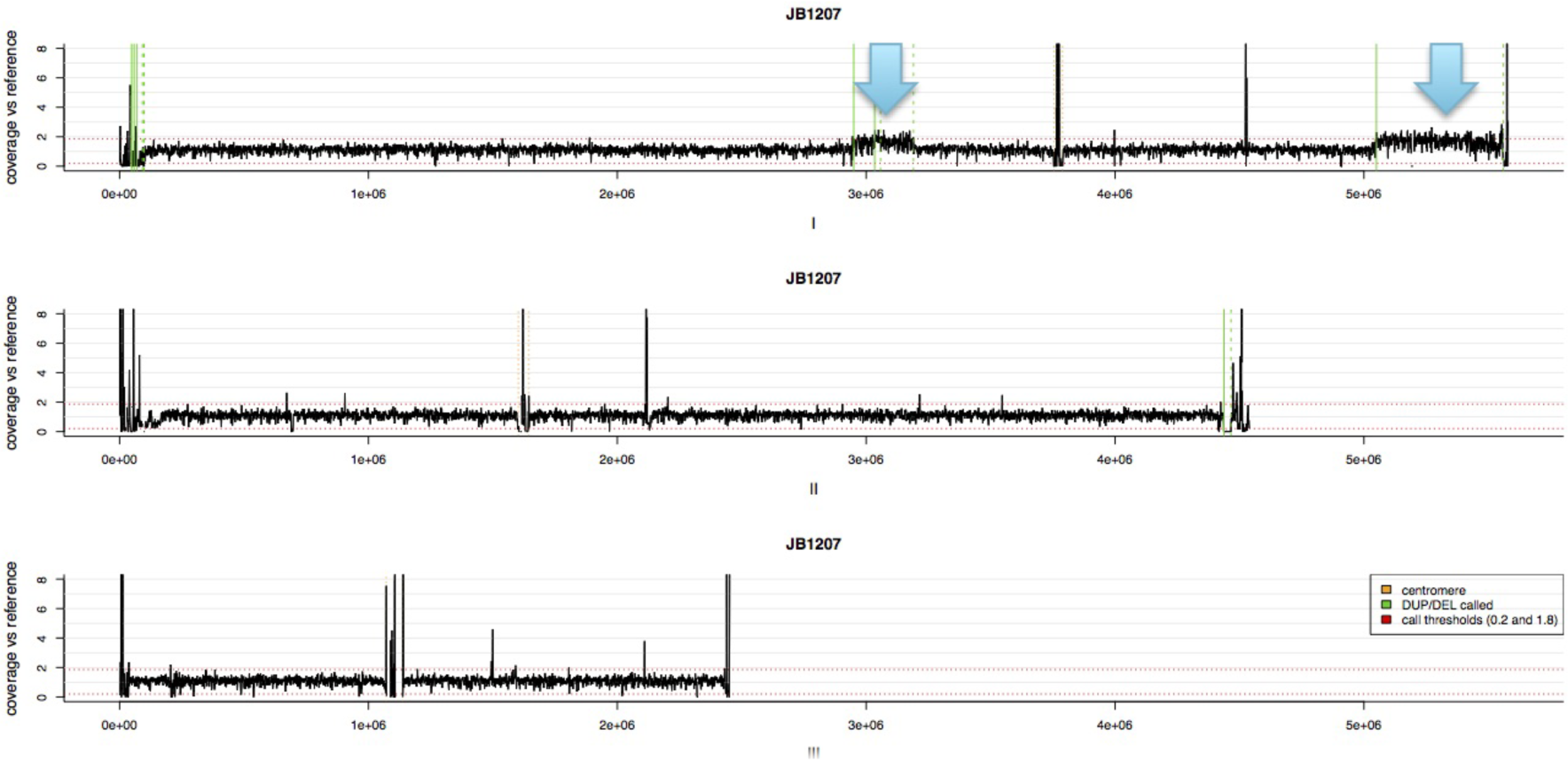
Chromosome-scale read coverage plots for three chromosomes of strain JB1207. Coverage is calculated relative to the reference strain (JB22 in our collection). Two large duplications that did not satisfy the criteria used to detect CNVs with cn.MOPs are indicated with blue arrows (DUP.I:2950001..3190000, 240kb and DUP.I:5050001..5560000, 510kb).

